# Statistical Inference of Enhancer-Gene Networks Reveals Pivotal Role of T-bet Expression Intensity for T Helper Cell Fate

**DOI:** 10.1101/2022.11.23.517729

**Authors:** Christoph Kommer, Qin Zhang, Ahmed N. Hegazy, Max Löhning, Thomas Höfer

## Abstract

Mammalian genomes harbor many more enhancers than genes, which greatly complicates the elucidation of cell-state-specific regulatory networks. Here, we developed a computational framework for learning enhancer-based gene networks from joint data on enhancer activity and transcript abundance. Dissecting the developmental plasticity of T helper (Th) cells with this approach, we uncovered a highly connected enhancer-gene network that supports graded Th-cell differentiation states, rather than mutual exclusivity of type-1 and type-2 immunity. Machine learning identifies a small number of regulatory enhancer types as network hubs. Hub enhancers in Th1 cells integrate as inputs the expression level of the master-regulator transcription factor, T-bet, and STAT signals governed by the cytokine environment. The quantitative balance between cell-intrinsic T-bet, driving phenotypic stability, and environmental cues enabling plasticity explains the heterogeneous reprogramming capacities of individual Th1 cells differentiating during natural infections *in vivo*. Moreover, we provide a framework for elucidating genome-scale regulatory networks based on enhancer activity.

## INTRODUCTION

The mammalian immune response is a paradigm for the functional differentiation of polarized lymphocytes, providing effective defense to diverse classes of pathogens. Type-1 immunity, directed against pathogen-infected cells, and type-2 immunity, aiming at extracellular pathogens, have long been recognized as major branches (Spellberg and Edwards, 2001; Yamaguchi et al., 2015). At the transcriptional level, the fate decisions of immune cells were shown to be governed by the expression of master regulators. Specifically, T-bet (encoded by Tbx21) and GATA-3 are the master regulators, respectively, of T-helper type-1 (Th1) and type-2 (Th2) lymphocytes (Murphy and Reiner, 2002). Cell-autonomous autoactivation loops stabilize both GATA-3 and T-bet expression, and there are numerous inhibitory interactions between Th1- and Th2-inducing molecular pathways (Ouyang et al., 2000; Mullen et al., 2001; Afkarian et al., 2002; Peine et al., 2013; Kanno et al., 2012; Löhning et al., 2002). While these findings have given rise to a model positing stability and mutual exclusion of Th1 and Th2 cell states, much of the underlying experimental work focused on experimental conditions geared towards pure Th1 or Th2 cell responses. Subsequently, however, data have been accumulated that show graded cell states along the Th1-Th2 axis that are not transitory but long-term stable (Hegazy et al., 2010; Antebi et al., 2013; Huang, 2013; Peine et al., 2013). This fine-tuning is vital for balancing effective immune responses while limiting immunopathology. Mechanistically, however, the remarkable combination of fine-tuning and stability of Th cell states has been surprising, as conventional models posit discrete and mutual exclusive Th cell types demarcated by mutual inhibition of master regulators and subseuent epigenetic imprinting (Murphy and Reiner, 2002; Huang, 2013).

Ultimately, gene-regulatory and epigentic networks encode the stability and plasticity of immune cell states. Seminal work has shown that key environmental sensors for Th cell activation and differentiation, the JAK-STAT4 and STAT6 pathways, shape the enhancer landscape of T helper cells prior to the action of T-bet and GATA-3 (Vahedi et al., 2012). This is particularly interesting as the STAT1 and STAT6 pathways driving Th1 and Th2 differentiation are also pivotal in determining the fate of type-1 and type-2 macrophages (Gordon and Martinez, 2010). This work emphasizes the long-standing question of whether cytokine signal transducers can stably imprint Th cell fates on their own, or whether transcriptional master regulators are required.

Classically, gene-regulatory networks are defined by connecting genes (the nodes of these networks) by activating or inhibitory regulatory interactions (the network edges). However, we now know that mammalian genes typically receive input from several enhancers, allowing for complex logic that is not captured by binary gene-to-gene connections (Yue et al., 2014). Experimentally, much progress has been made in probing the activity state of enhancers and in cataloging the transcription factors that bind to enhancers in a genome-wide manner (Spitz and Furlong, 2012; Andersson and Sandelin, 2019). Integrating these data into gene-enhancer networks requires the comprehensive identification of the genes that are regulated by all the relevant enhancers. Given the enormous number of enhancers, which is about an order of magnitude larger than the number of genes (Yue et al., 2014), experimental data remain incomplete, and faithful computational inference approaches for enhancers and their target genes are being called for. Initially, such approaches were based on gene-enhancer proximity. This criterion does not reflect the pervasive finding that important enhancers can be located hundreds of kbp away from the start site of the gene they regulate. A more promising approach consists in exploiting the co-variation of enhancer activity and gene expression, which was recently developed to derive disease-specific gene-enhancer networks by exploiting inter-individual variability in these variables (Reyes-Palomares et al., 2020).

Here, we develop a direct perturbative approach to infer the gene-enhancer network governing the plasticity of Th1 lymphocytes along the axis towards type-2 immunity. To comprehend the interplay of the master-regulator transcription factor T-bet with environmental sensing through STATs, we combine dosing of T-bet expression quantities in primary T cells with exposing these cells to different cytokine environments. From these experiments, we learn the gene-enhancer network underlying Th-cell differentiation states, interrogate its functioning by machine learning, and apply our findings to rationalize the behavior of *in vivo* generated Th1 cells towards opposing differentiation signals (Hegazy et al., 2022).

## RESULTS

### Differentiation and reprogramming of helper T cells globally alters key histone modifications

To study global epigenetic changes during differentiation and reprogramming of T helper cells, we used Th1 cells primed in vivo under strong Th1-polarizing conditions induced in mice by infection with lymphocytic choriomeningitis virus (LCMV). Naïve TCR-transgenic LCMV-specific CD4+ T cells were adoptively transferred into C57BL/6 mice (WT mice), which were then infected with LCMV (Figure S1). Subsequently, we sorted in vivo-primed LCMV-specific Th1 cells from the infected hosts and cultured them under neutral cytokine conditions to perpetuate the Th1 phenotype, or under Th2 cytokine conditions to re-program them into cells combining functional properties of Th1 and Th2 cells, termed Th1+2 cells (Figure 1A; Figure S1; cf. (Hegazy et al., 2022)). As control, we directly differentiated naive Th cells into Th2 cells *ex vivo*. While T-bet (*Tbx21*) had initially been identified as the master-regulator transcription factor for Th1 cell identity (Szabo et al., 2000), subsequent work emphasized the pivotal role of the cytokine-activated transcription factors STAT1 and STAT4, rather than T-bet, in shaping the epigenetic landscape of Th1 cells (Vahedi et al., 2012). To address the role of T-bet in regulating Th1 cell plasticity, we performed all experiments with cells harboring two (wildtype, WT), one (heterozygous, HET), or no (knockout, KO) functional *Tbx21* alleles; these genotypes result in corresponding high, intermediate, or absent capacity to express the T-bet protein, respectively (Hegazy et al., 2022). Combining neutral or Th2-polarizing cytokine environments and *Tbx21* doses, we obtained six conditions for examining epigenetic stability versus plasticity of Th1 cells, together with two controls, naive and Th2 cells (Figure 1B). We performed ChlP-seq for key permissive (H3K4me1, H3K4me3, and H3K27ac) and repressive (H3K9me3 and H3K27me3) histone modifications for all conditions, achieving high reproducibility of biological replicates (Figure S2).

**Figure 1:**
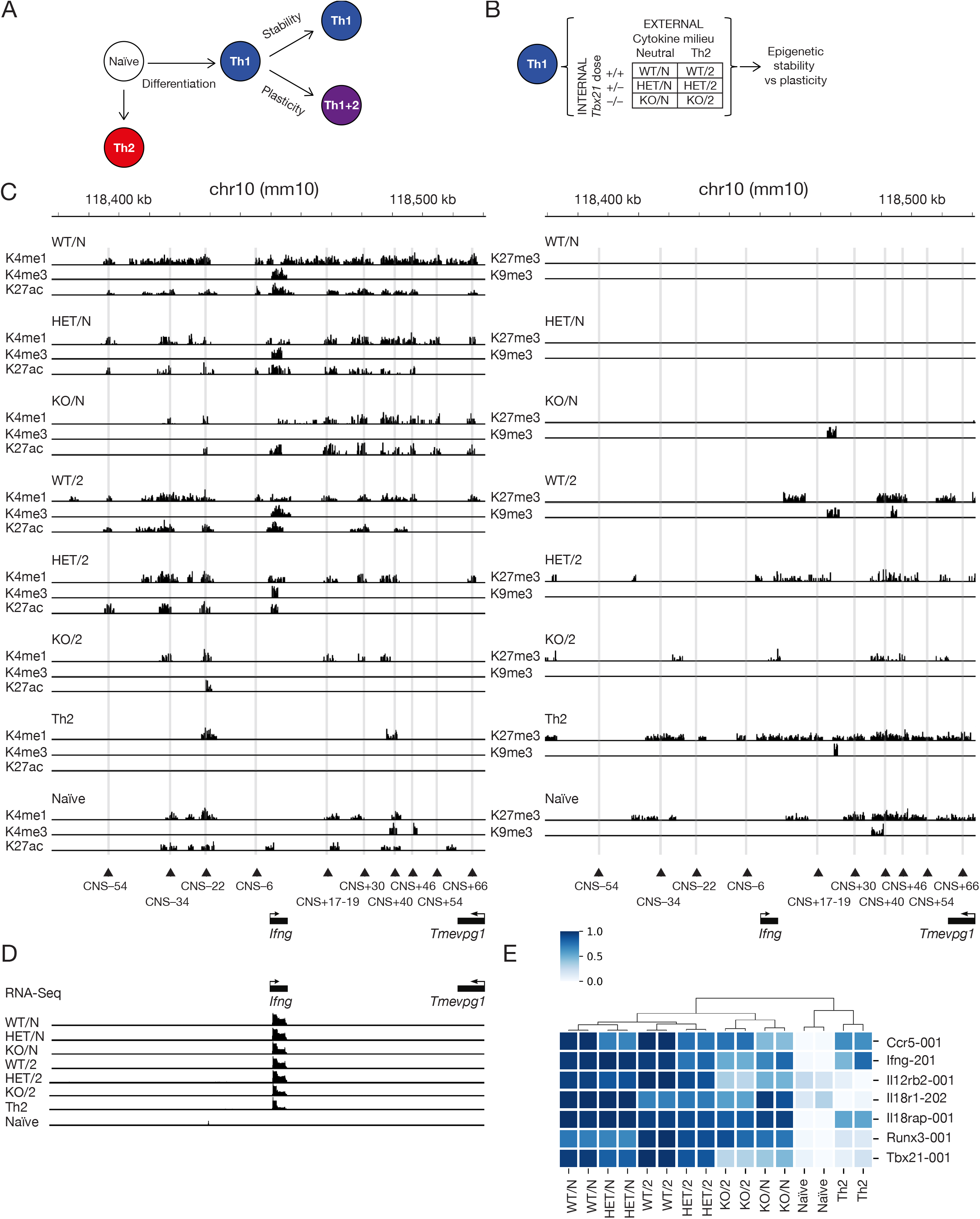
Differentiation and reprogramming of helper T cells globally alters key histone modifications. (A) Naïve cells were either differentiated into Th1 or Th2 cells. The Th1 cells were kept under Th1/neutral cytokine or Th2 cytokine conditions to obtain stability as well as plasticity features. (B) Th1 cells were obtained with wild-type (WT), heterozygous (HET) and knock-out (KO) *Tbx21* under neutral and Th2 cytokine conditions. Subsequently we assessed differences in epigenetic stability and plasticity. (C) Significant histone modification peaks at the signature Th1 cytokine locus of *Ifng*. On the left side we listed all permissive marks for one replicate of all experimental conditions; on the right hand side we listed the same for the repressive marks. The grey vertical bars mark conserved nucleotide sequences representing known enhancers. (D) Raw RNA-Seq read data on log scale in direct comparison to the histone modification peaks shown before. (E) Selection of notable Th1 gene loci showcasing similar transcriptional behaviour over experimental conditions and replicates. Hierarchical clustering over conditions shows the evident T-bet sensitivity and distinction from naïve and Th2 conditions.

To illustrate key features of the data, we focus on the signature Th1 gene locus *Ifng*, the enhancers of which have been annotated and functionally studied in considerable detail (Balasubramani et al., 2010). Th1 cell differentiation during virus infection induced permissive histone marks (Figure 1C, left panel) and eradicated repressive marks (Figure 1C, right panel) overall. The induction of permissive marks was weaker with lower T-bet dose while the removal of repressive marks was not affected by T-bet dose (except for one position, where H3K9me3 appeared in *Tbx21* cells (KO/N)). By contrast, exposure of Th1 cells to Th2 cytokine conditions caused both a reduction of permissive and an induction of repressive histone marks. This result is consistent with direct Th2 differentiation reducing permissive marks found already in naive Th cells and increasing repressive marks. Constitutive *Ifng* transcription was strongest in *Tbx21* Th1 cells and declined with decreasing T-bet dose and exposure to Th2 cytokine conditions (Figure 1D). These effects were characteristic of other associated Th1 gene loci as well (Figure 1E). Taken together, these findings suggest distinct effects of T-bet dose and cytokine stimuli on the epigenetic plasticity of Th1 cells.

### Enhancer activities respond to T-bet dose or/and cytokine stimuli

To identify enhancers and other regulatory elements from the patterns of combined histone modifications, we used a Hidden Markov Model implemented in ChromHMM (Ernst and Kellis, 2010). Testing for different potential numbers of chromatin states, the information in the classification plateaued at 16 states (Figure 2A; Figure S3). States 1 and 2 are associated with poised and active enhancer regions respectively (marked, respectively, by H3K4me1 and by H3K4me1 and H3K27ac combined). In contrast to a previously proposed approach (Reyes-Palomares et al., 2020), we additionally account for the repressive mark H3K27me3 in the definition of enhancer activity states, finding bivalent enhancer States 4 and 5 (H3K4me1 or H3K4me1/H3K27ac combined with H3K27me3) as well as repressed State 3 (H3K27me3 only). Finally, chromatin domains with the silencing mark H3K9me3 were rare in loci with Th cell-specific genes.

**Figure 2:**
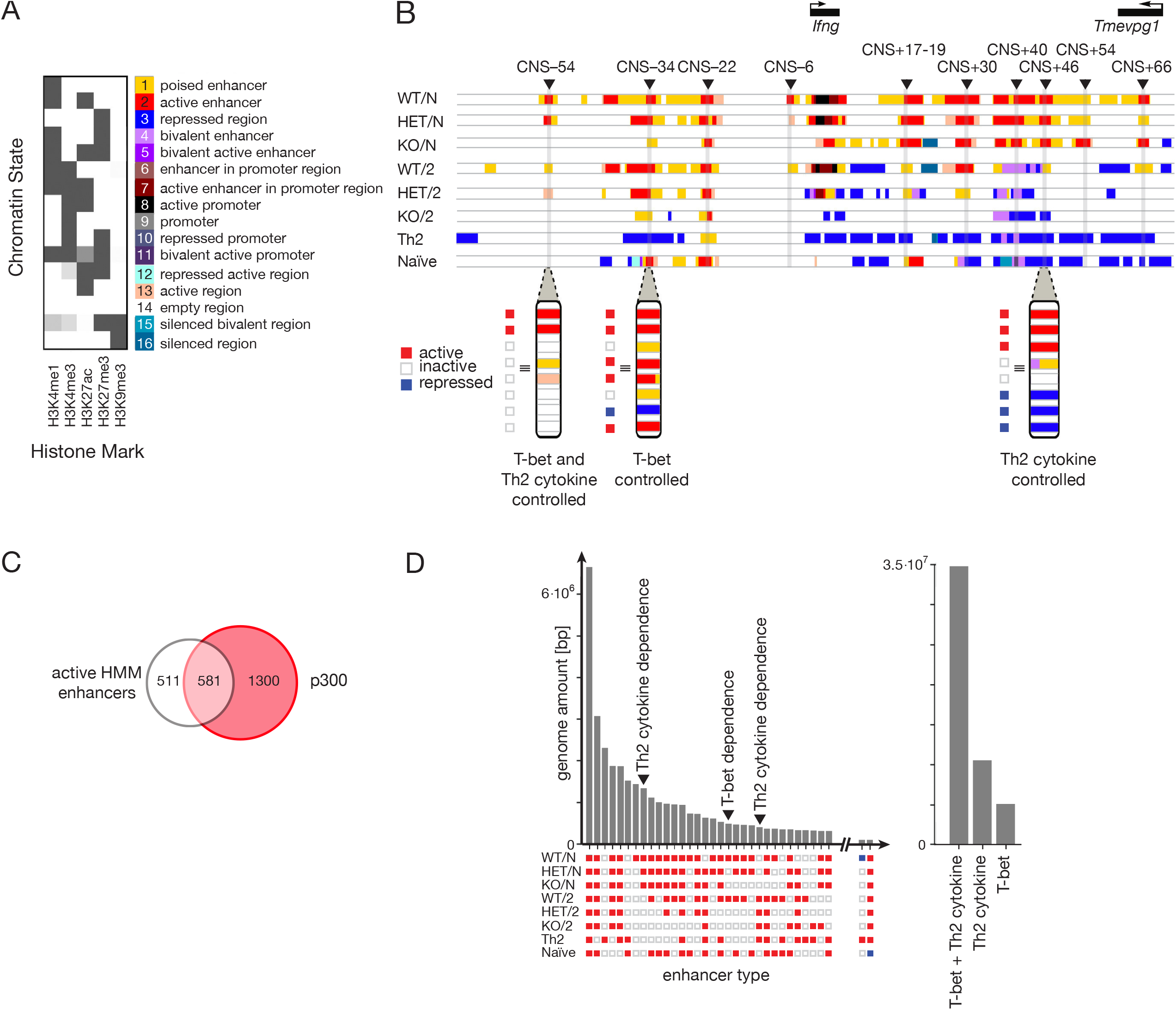
Enhancer activities respond to T-bet dose or/and cytokine stimuli based on a Hidden Markov Model. (A) Best fit Hidden Markov Model (HMM) from ChromHMM consisting of 16 colour-coded chromatin states. The heatmap shows the emission probabilities of finding a certain histone mark in a certain labelled state. Most notably we found enhancer states, repressed states, promoter states and bivalent states of which States 1, 2 and 3 are of utmost interest. (B) Full chromatin state landscape at the *Ifng* locus based on the inferred HMM for all experimental conditions. We found three main types of enhancers – Tbet controlled, Th2 cytokine controlled and a mixture of both – when considering an enhancer either to be active, inactive or repressed. Chromatin state colours are the same as in (A). (C) p300 binding overlap with active HMM enhancers in either Th1 or Th2 conditions. (D) Frequency distribution in terms of the genome amount in bp of regulatory enhancer types around Th1 and Th2 genes (left). Enhancer types consist of active (red), inactive (white) and repressed (blue) enhancers depending on the experimental condition. The summary in terms of T-bet and Th2 cytokine dependence is shown on the right.

We validated the enhancer classification for the *Ifng* locus, identifying all previously described enhancers (designated as conserved non-coding sequences, CNS; Figure 2B) (Balasubramani et al., 2010). All these enhancers were classified as active in *Tbx21^+/+^* Th1 (WT/N) cells, with the exception of CNS+54 where H3K27ac did not rise above the significance threshold. To check to what extent the enhancers identified via histone modifications overlap with an alternative enhancer definition by p300 binding (Vahedi et al., 2012), we analyzed all active enhancers in topologically associating chromatin domains (TADs) containing Th1-specific genes (Table S1). Of these 1092 active enhancers, 53% had a p300 binding peak in *in vivo–* differentiated Th1 cells (Figure 2C), representing a large and highly significant overlap.

All enhancers in the *Ifng* locus were active only in a subset of conditions, becoming inactive or even repressed in other conditions. Three characteristic patterns of activity changes emerged, exemplified by CNS+46, CNS-34, and CNS-54 (Figure 2B). The enhancer at CNS+46 was repressed in naive cells, active in Th1 cells of all T-bet doses in the absence of Th2 cytokines, and inactive or repressed in the presence of Th2 cytokines; therefore, its activity was primarily altered by Th2 cytokine stimuli. Conversely, the enhancer at CNS-34 became inactive in Th1 cells only when T-bet was not present but did not respond to the presence of Th2 cytokines; hence its activity was controlled by T-bet. The CNS-54 enhancer represented a mixed type, being responsive both to T-bet dose and Th2 cytokines. These examples suggest condensing the finely resolved characterization of enhancer activity through ChromHMM to a coarse-grained classification of an enhancer as either active, inactive (e.g., poised or bivalent), or repressed. In this way, we characterize each enhancer by its activity pattern across the experimental conditions. All enhancers that share a given activity pattern are considered an *enhancer type*, characterized by a specific mode of regulation by T-bet dose and cytokine environment. We computed the abundance of all occurring enhancer types across the genome, as measured by the total genome length they occupy. Enhancers active in all conditions were the single most abundant type, and these are most likely linked to fundamental aspects of cell integrity and T cell identity (Figure 2D). However, the sum of all differentially regulated enhancers exceeded the constitutively active ones by about five-fold, with enhancer types responding to both T-bet dose and Th2 cytokines being the most frequent (Figure 2D). Thus, the reprogramming of Th1 cells is associated with global changes in the enhancer landscape.

### A multivariate enhancer activity score predicts enhancer-target gene associations

Next, we asked whether enhancer activity states correlate with gene transcription. To this end, we computed the correlations between the amount of histone modifications that characterize enhancer activity in the ChromHMM classification, namely H3K4me1, H3K27ac, and H3K27me3, and the transcript copy number, focusing on known enhancer-gene associations. As expected, we found strong positive correlations between H3K27ac and transcript number while H3K4me1 and H3K27me3 were also – positively and negatively, respectively, – correlated (Figure 3A). Hence, we define the following enhancer activity score:

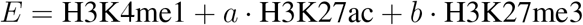

for a given enhancer and experimental condition (Figure 3B). The weights a and b quantify the relative influence of each mark on enhancer activity. To predict unknown enhancer-gene interactions, we determined these weights by maximizing the Pearson correlation between enhancer activity score and transcript copy number for a training set of 14 experimentally validated enhancers of Th1 and Th2 genes (Table S2; Figure S4 A,B), finding *a* = 1.2 ± 0.7 and *b* = 2.8 ± 1.1. Thus, H3K4me1 and H3K27ac have comparable weight in defining an active enhancer and, hence, combining these two marks leads to more robust recognition of active enhancers than using H3K27ac alone. Moreover, the large negative weight of H3K27me3 underscores the importance of this modification for predicting enhancer activity. The presence of H3K27me3 (i.e., an enhancer being located in a repressive chromatin domain) can fully offset the effects of activating enhancer modifications.

**Figure 3:**
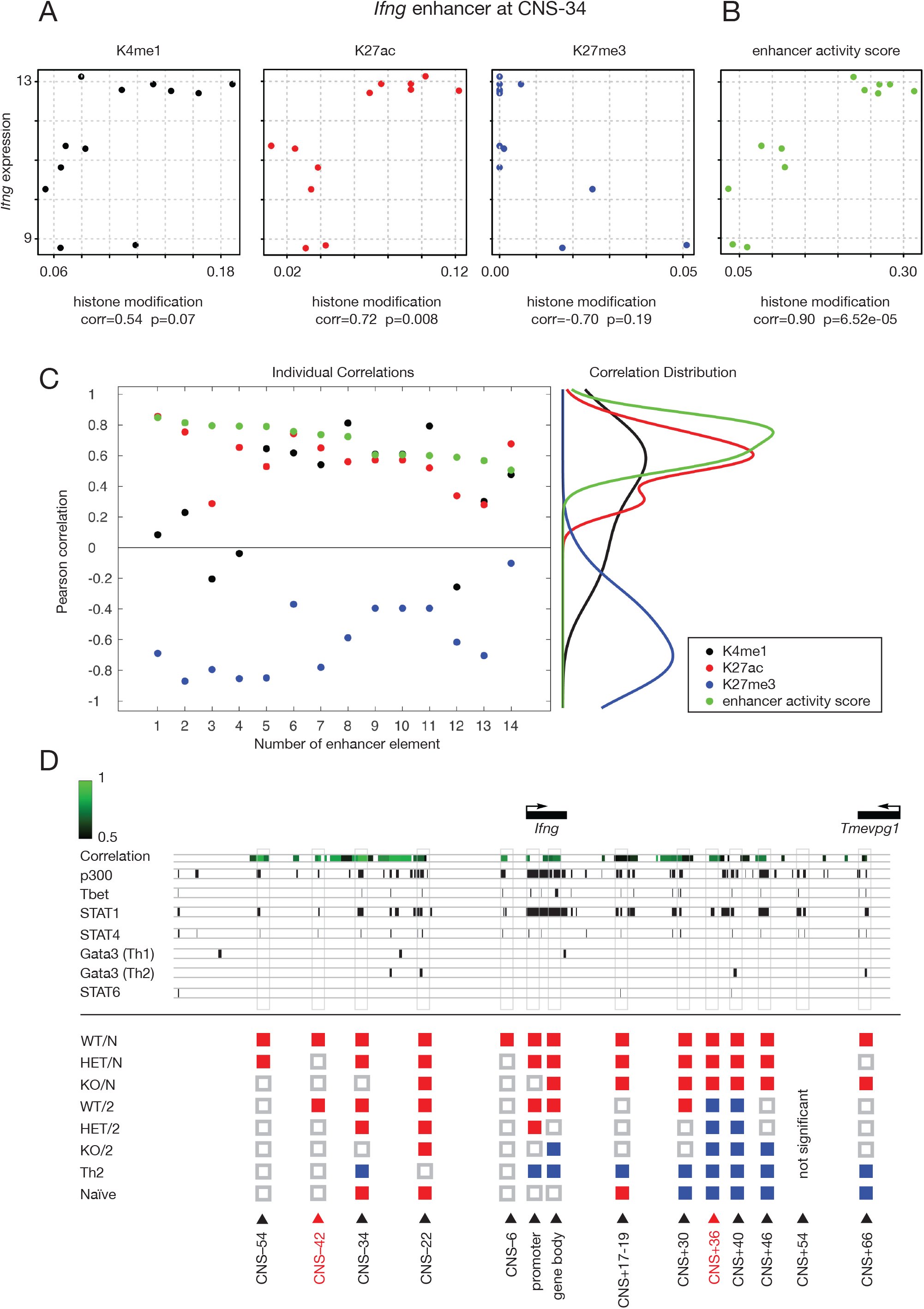
A multivariate enhancer activity score predicts enhancer-target gene associations at *Ifng*. (A) Correlations of H3K4me1, H3K27ac, and H3K27me3 with *Ifng* transcript abundance at the enhancer element CNS-34 upstream of *Ifng*. (B) Correlation of the parametrized enhancer activity score at CNS-34 with *Ifng* transcript abundance. (C) Individual Pearson correlations for the enhancer learning sample (numbering see Table S1) of H3K4me1, H3K27ac, H3K27me3 as well as the enhancer activity score with the respective gene transcript abundances. The distributions of the respective correlations are shown on the right. (D) Prediction of significant correlations according to our model at the *Ifng* locus. Significant Pearson correlations are shown in shades of green accompanied by transcription factor bindings (Nakayamada et al., 2011; Vahedi et al., 2012; Wei et al., 2010; Wei et al., 2011) labelled by the respective enhancer types at these positions (active: red; inactive: white; repressed: blue). Newly predicted CNS enhancer sites are marked in red.

Across the learning sample, the enhancer activity score performed significantly better than any single histone mark in correlating with transcript abundance (Figure 3C). This result indicates that this score can be used to predict target genes of enhancers. Indeed, applying it to the entire *Ifng* locus with a statistical significance test, we recovered all but one (CNS+54 which was just below significance) of the ten described enhancers, including three enhancers that were not part of the learning sample. Moreover, we predict two new enhancer elements. Checking for sequence conservation and transcription factor binding, we found that these putative new enhancers correspond to conserved non-coding sequences (CNS-42 and CNS+36), bind STAT1 and, in the case of CNS-42, also p300 and STAT4 (Figure 3D). Taken together, these findings indicate that the enhancer activity score reliably maps enhancers to their target gene.

### Partial correlations identify unique enhancer-gene associations in topologically associating domains

Topologically associating domains (TADs), typically ranging between 200 kB and 2.5 Mb (Gong et al., 2018), are a biologically meaningful unit to search for enhancer-transcript correlations (Barrington et al., 2019). Most TADs contain several genes, posing the question of whether enhancers map uniquely to individual genes or activate all genes within a TAD. A prominent example is the Th2 cytokine locus which resides within one TAD and contains the *Il4, Il5*, and *Il13* genes as well as three genes unrelated to adaptive immunity (Figure 4A) (Ansel et al., 2006). Naively computing correlations of enhancer activity score and transcript abundance across this locus would predict extensive overlap of enhancers for the three cytokine genes as well as the non-immune genes *Rad50* and *Sept8* (Figure 4B).

**Figure 4:**
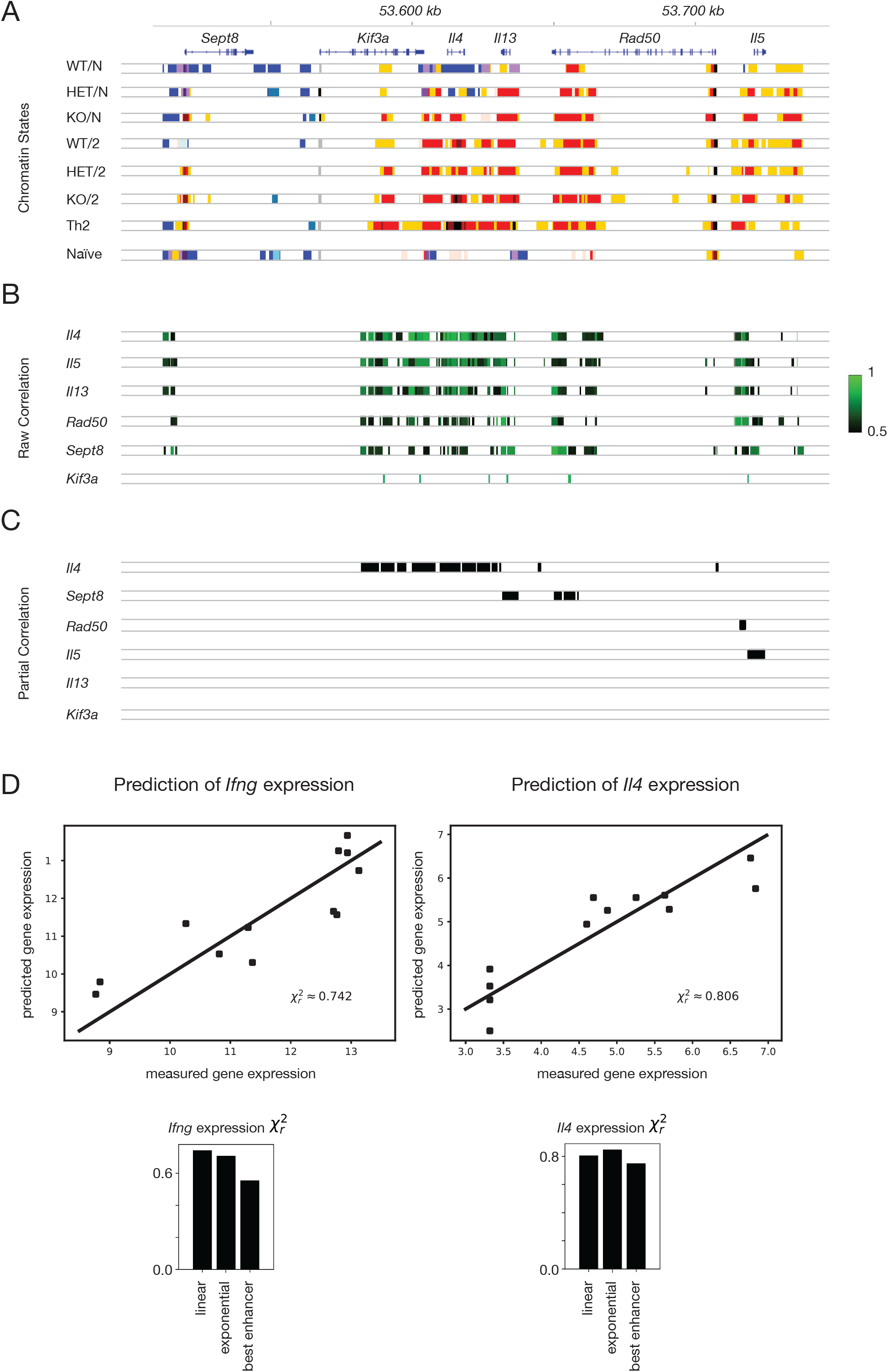
Partial correlations identify unique enhancer-gene associations in topologically associating domains at the Th2 cytokine locus and are able to predict gene expression. (A) Central part of the Th2 cytokine locus on chromosome 11 with *Sept8, Kif3a, Il4, Il13, Rad50* and *Il5* with the according HMM state classification. (B) Significant zero-order Pearson correlations of each enhancer activity score segment with each gene in the locus in shades of green according to our model. We observe substantial coregulation in this case. (C) Significant partial correlations of the enhancer activity score segments with each gene removing spurious correlations with all remaining genes respectively. (D) Prediction of *Ifng-expression* (left) and Il4-expression (right) by a linear model of enhancer activity weighted by their respective correlations (top) based on a reduced χ^2^ statistic. Comparison with models including non-linear enhancer interactions (exponential) and only choosing the highest correlating enhancer (bottom).

However, correlations of a given enhancer with multiple transcripts may be spurious if there are additional, confounding factors that cause correlations between transcripts, yet only one transcript (or a subset) is actually controlled by the enhancer. We found that partial correlations (Guo et al., 2017) better measure the association between transcript abundance and enhancer activity, as they strongly reduce the effect of confounding factors. Applying partial correlations to the data on the Th2 cytokine locus identified non-overlapping gene-specific enhancers for *Il4*, *Il5*, *Rad50*, and *Sept8* (Figure 2C), with the high density of regulatory elements for *Il4* being consistent with previous work (Ansel et al., 2006). Taken together, the enhancer activity score combined with partial correlation analysis can be used to identify gene-specific enhancers and moreover predict gene expression.

Next, we asked if gene expression can be predicted from enhancer activity. We first tested a model where all enhancers contribute independently and the impact of each individual enhancer is given by its enhancer activity score correlation. This model predicted the expression of the signature cytokines *Ifng* and *Il4*, based on gene-specific enhancers identified by partial correlations (Figure 4D). A model with cooperative enhancer interactions had similar predictive power, whereas a dominant-enhancer model, with the most strongly correlating enhancer overriding the effect of the other enhancers on target gene transcription, performed substantially worse for both genes. Hence, the level of gene transcription is a joint function of all the enhancers of the respective gene.

### Enhancer activity patterns distinguish cell-type-specific genes

We applied our method to systematically infer enhancer-gene associations to identify the enhancers regulating a core set of 96 transcripts differentially expressed in Th1 versus Th2 cells as derived from our RNA-Seq analyses and as shown in (Wei et al., 2011). We identified 6168 enhancers, the vast majority of which had not previously been mapped to target genes. Moreover, known enhancers were refined by this analysis; in particular, an apparent *Tbx21* superenhancer was decomposed in a set of five discrete enhancer elements (Figure S5). Next, we identified the regulatory enhancer types based on the ternary classification of active, inactive, and repressed enhancers. The 6168 Th1/Th2-related enhancers belonged to 670 regulatory enhancers types, suggesting that the focus on enhancer types instead of individual enhancers will considerably simplify the analysis of the gene-regulatory logic underlying Th1 cell plasticity. This is even more so the case, as we will show now that only a subset of these enhancer types plays a prominent role.

To understand which enhancer types are key regulators of Th1 cell plasticity, we asked whether and to what extent a given type predicts Th1 versus Th2 differentiation. To this end, we used a standard tool of machine learning, a tree-based supervised learning algorithm (Extremely Randomized Trees; (Geurts et al., 2006)), to classify genes as Th1-specific or Th2-specific based on their associated enhancers. This approach identified key Th1-promoting and Th2-promoting enhancer types (Figure 5A). The four highest ranked Th1-promoting enhancer types were sensitive to T-bet dose, as they were inactive in *Tbx21^-/-^* cells and, except for two conditions, also in *Tbx21^+/-^* cells (Figure 5A left). The majority of these enhancer types was dependent on both T-bet dose and cytokine signals; the second largest group was exclusively T-bet dependent, whereas enhancer types that responded exclusively to Th2 cytokine conditions were rare (Figure 5B left). These findings show that the presence of T-bet, and for enhancers inactive in *Tbx21^+/-^* cells also the T-bet dose, is a key a determinant of Th1-specific enhancer activity.

**Figure 5:**
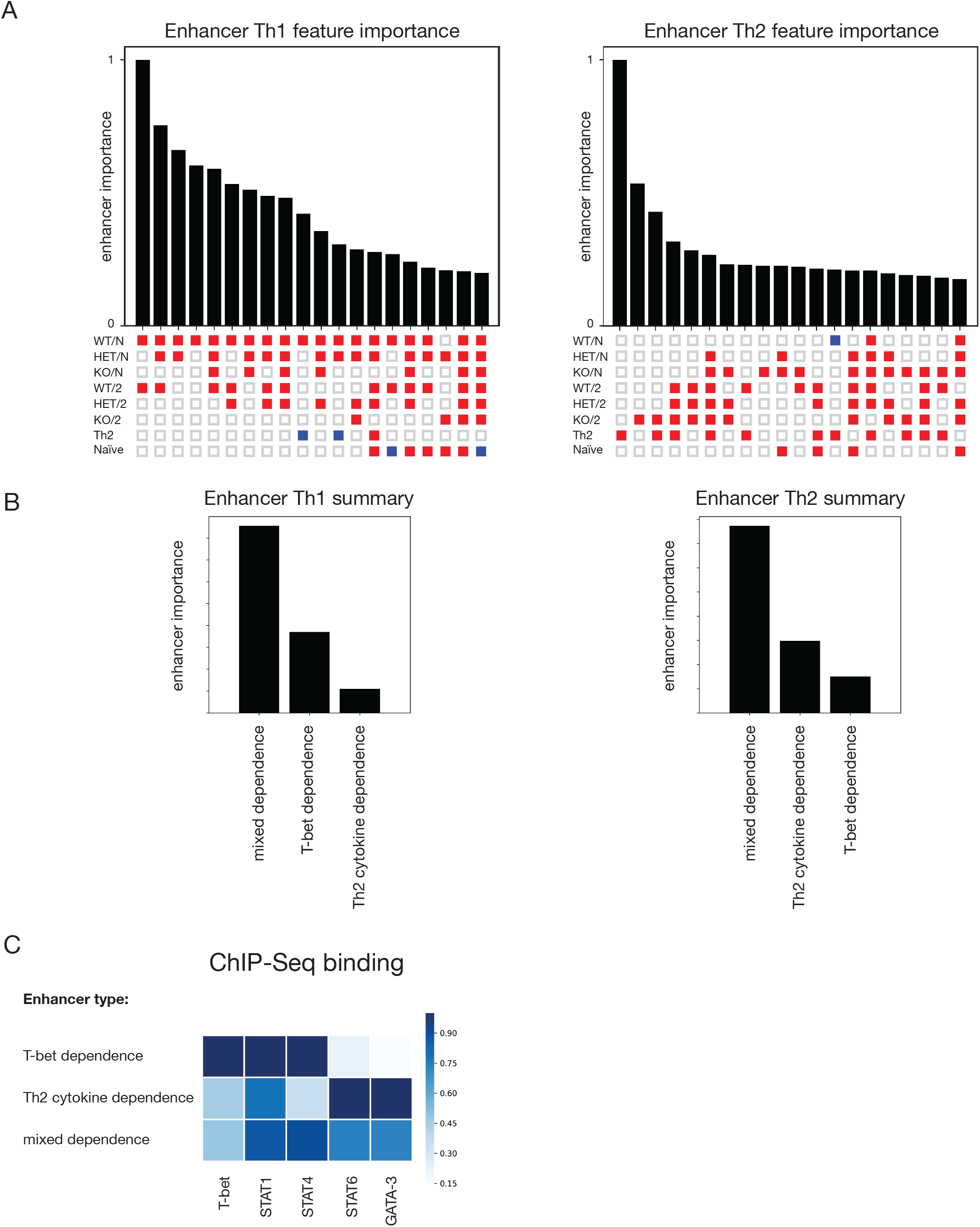
Enhancer activity patterns distinguish cell-type-specific genes and exhibit distinct transcription factor regulation. (A) Importance ranking of the top 20 regulatory enhancer types (active: red; inactive: white; repressed: blue) belonging to Th1-specific genes (left) as well as to Th2-specific genes (right). (B) Summary of the importance assigned to the three main regulation categories of enhancer types mapped to Th1-specific genes (left) and Th2-specific genes (right). (C) TF regulation of the three main enhancer types around Th1 and Th2 genes. The respective TF weight is shown in shades of blue.

The highest ranked Th2-promoting enhancer type was active only in Th2 conditions and hence was activated exclusively by direct Th2 differentiation of naïve T cells; the enhancers belonging to this type are therefore not susceptible to Th1→Th2 reprogramming (Figure 5A right). However, a large set of enhancers was active in Th2 cells and was switched on during reprogramming; the majority of these reprogrammed enhancers were activated under conditions of low or absent T-bet (Figure 5A right). Indeed, most Th2-promoting enhancers responded to both T-bet (negatively) and Th2 cytokine signals (positively), whereas only a minority of Th2-promoting enhancers were exclusively responsive to Th2 cytokines (Figure 5B right). Thus, T-bet also acts as a key gatekeeper of Th1→Th2 reprogramming (Hegazy et al., 2022).

For each enhancer type, we analyzed binding of transcription factors to identify common modes of regulation for enhancers of this type. Aggregating these data across all Th1-promoting and Th2-promoting enhancers we found that the three main enhancer categories - controlled by T-bet and Th2 cytokine conditions, controlled by T-bet only, and controlled by Th2 cytokines only - had characteristic modes of regulation (Figure 5C). Exclusively Th2-cytokine-dependent enhancers were bound mostly by the signal transducer downstream of IL-4, STAT6, and the STAT6-induced master-regulator transcription factor GATA-3, whereas T-bet, STAT1 or STAT4 binding were much less frequent (note that STAT1 has generally many more genomic binding events in ChIP-Seq data than the other transcription factors; (Vahedi et al., 2012)). An inverse pattern was observed at the T-bet-controlled enhancers that hardly bound STAT6 and GATA-3 but instead T-bet, mostly together with the T-bet inducing factors STAT1 and STAT4. The largest group of enhancers, controlled by T-bet and Th2 cytokines, bound both the Th1- and Th2-associated STATs whereas T-bet binding was reduced compared to exclusively T-bet-controlled enhancers.

In conclusion, we inferred subtype specificity of Th cell enhancers in a genome-wide manner, and find an underlying regulatory logic in differential transcription factor binding. The vast majority of Th1-promoting enhancers as well as most enhancers relevant for Th1→Th2 reprogramming are T-bet sensitive

### Enhancer-gene networks reveal condition-specific topologies

To elucidate the regulatory logic underlying Th1-cell plasticity, we integrated the information on Th-subtype-specific enhancers, their regulating transcription factors, and their target genes by building bipartite enhancer-gene networks (Figure 6A). We obtained a densely connected network. Nevertheless, a force-directed layout of the network revealed clear separation of Th1- and Th2-specific regions, joined by a smaller intermediate domain (Figure 6B). Genes that are specific for Th1 or Th2 conditions lie within the respective parts of the network, as do the Th subtype-specific enhancer types (Figure S6). The network includes both positive interactions, where binding of a TF activates an enhancer and, in turn, its target gene(s), and negative interactions where the presence of a transcription factor inhibits enhancer and target gene(s) (Figure S6). The network layout with distinct Th1-specific and Th2-specific enhancer-gene regions implies coherence of the respective gene-regulatory programs and allows for graphic inspection of the Th cell polarization state.

**Figure 6:**
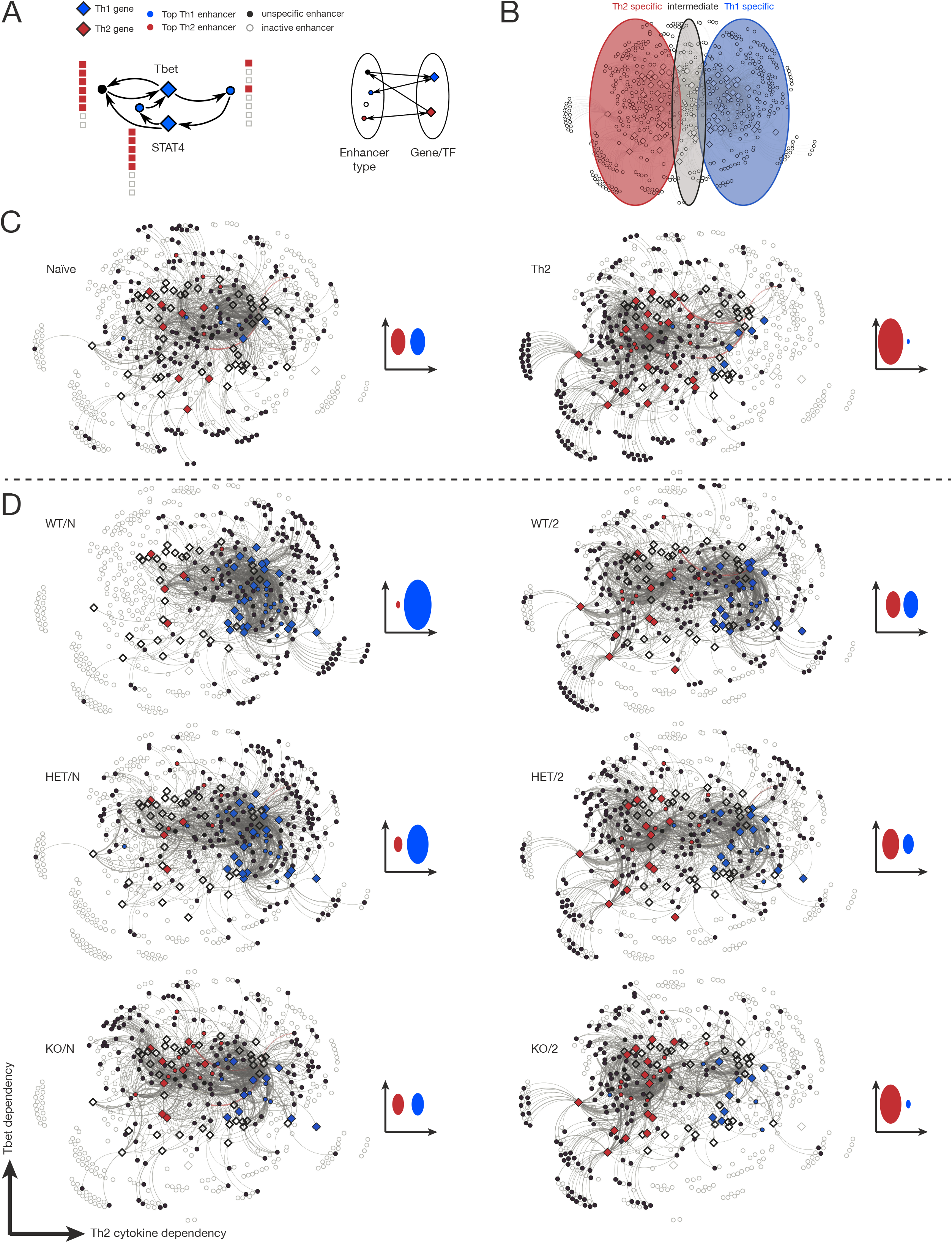
Enhancer-gene networks reveal condition-specific topologies in multiplex net-works. (A) Network architecture example for T-bet and Stat4 and actual binding of enhancer types (order of experimental conditions as before) and vice versa (left). The bipartite bidirectional structure is shown as enhancer types are always mediated by genes and TFs and vice versa (right). (B) Node positions of the full enhancer-gene network according to a force-directed depiction. The left side turns out to contain Th2-specific nodes, while the right side contains Th1-specific nodes. The intermediate region contains a mixture of both as well as unspecific nodes. (C) Condition-specific multiplex networks for naïve and Th2 conditions. Inactive enhancers (summarized by enhancer types) as well as genes with a low or basal expression level are labelled in white; active enhancers and highly expressed genes are labeled in black. Th1 specific genes and enhancers are blue and Th2 specific ones are red. For each network a schematic distribution of enhancer-type and gene activity in the Th1- and Th2-specific part of the network is given. (D) Condition-specific multiplex networks for the remaining experimental conditions with particular focus on T-bet dose for Th1 and Th1→Th2.

The generic network of Figure 6B integrates over all experimental conditions and hence displays all potential regulatory interactions. To understand how these generic networks change in particular Th cell states, we generated condition-specific multiplex networks (Battiston et al., 2014) by only considering enhancer types that are active in a given experimental condition. Hence, depending on the condition of interest, many nodes and edges of the full network vanish, and many remaining nodes (enhancers and genes) switch activity. The naive Th network was unpolarized, with mostly subset-unspecific enhancer types (i.e., active in the naive condition) active in the network core and a homogeneous distribution of active enhancer types surrounding the core (Figure 6C, left). In stark contrast, in directly programmed Th2 cells enhancer activity switched over to the Th2 region and became depleted in the Th1 region (Figure 6C, right). The bona fide Th1 state represented the polarized counterpart, with the Th1-specific genes and corresponding enhancers active, and the Th2 region largely inactive (Figure 6D, WT/N). A decrease in T-bet dose alone caused only a moderate activation of enhancers on the Th2 side of the network (Figure 6D, HET/N), whereas *Tbx21* knockout caused more substantial activity of Th2-specific enhancers and considerable loss of Th1-specific enhancer activity (Figure 6D, KO/N). These data show that T-bet is required to activate Th1-specific enhancers and keep Th2-specific enhancers inactive genome-wide.

The reprogramming of Th1 cells under Th2 cytokine conditions also induced a shift to a more balanced distribution between Th1- and Th2-specific genes and enhancer types (Figure 6D, WT/2). Moreover, the effects of Th2 conditions and loss of T-bet became superimposed, to the point that *Tbx21* Th1-like cells reprogrammed under Th2 conditions exhibited a network topology strongly resembling the direct Th2 differentiation of naive T cells (Figure 6D, KO/2). In sum, T-bet dose and cytokine environment affect gradual - rather than binary - activity shifts in the enhancer-gene network.

### Differential regulation within networks is directed by specific enhancer types and is dependent on T-bet expression levels

To gain more detailed insight into Th1 cell reprogramming, we identified the specific enhancer types that differentially regulated reprogramming transitions. To this end, we computed up- and down-regulated enhancer-gene connections between pairs of condition-specific networks. The resulting differential networks (Figure 7A) show which genes lost input from their enhancers (due to enhancers becoming inactive, as indicated by downregulated edges in Figure 7A, orange lines) and which genes gained enhancer input (due to enhancers becoming active, as indicated by upregulated edges in Figure 7A, green lines). During Th1→Th2 reprogramming, we observed that Th1-specific genes (Figure 7A, blue diamonds) lost enhancer input whereas Th2-specific genes gained enhancer input (Figure 7A, red diamonds), with multiple enhancer types being involved in these gene switches. To rank these enhancer types, we searched for enhancer types that regulate many genes and also receive input from Th-cell-specific transcription factors. A robust procedure to identify this strong connectivity is the PageRank algorithm (Brin and Page, 1998), which we adapted to enhancer-gene networks. For Th1→Th2 reprogramming we identified a small number of pivotal enhancer subtypes regulating Th1- or Th2-specific genes (Figure 7B). Two of the three Th1-specific enhancer types are inhibited by Th2 cytokines and activated by T-bet (number 5 and 7), while the remaining Th1-specific enhancers (number 6) depend only on the presence of Th2 cytokines (Figure 7B). In a complementary manner, all four of the leading Th2-specific enhancer types were activated by Th2 cytokines (number 1 to 4), and two were additionally inhibited by T-bet.

**Figure 7:**
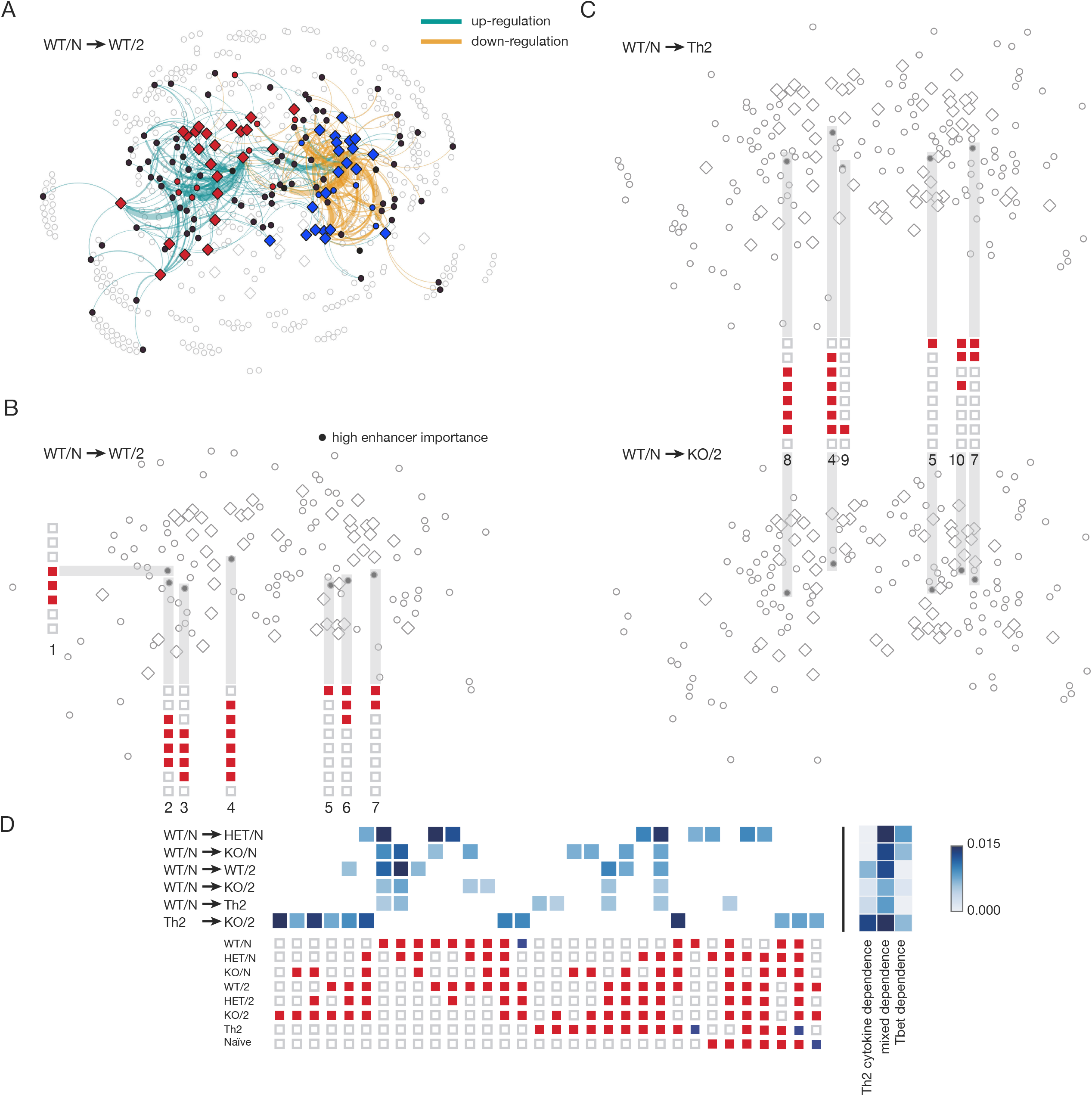
Differential regulation within networks is directed by specific enhancer types and is dependent on T-bet expression levels. (A) Differential regulation between the WT/N and the WT/2 network. Green line represents weighted up-regulated connections and yellow lines show the down-regulated connections. (B) Most important drivers in differential regulation of the WT/2 versus the WT/N network according to a random walk model. (C) Comparison of main drivers in differential regulation of WT/N→Th2 and WT/N→KO2 exhibiting large similarities. (D) Analysis of most important enhancer types in differential regulation of multiple pairs of experimental conditions. We show the impact of individual enhancer types (left) as well as the importances summarized in the three main classes of enhancer activity regulation (right).

During Th1→Th2 reprogramming of wild-type cells, T-bet levels decline only moderately; Th1+2 cells maintain much higher T-bet levels than Th2 cells (Hegazy et al., 2022). To evaluate specifically the effect of T-bet dose, we computed the differential network for the *Gedanken-experiment* of adding Th2 cytokines and knocking out *Tbx21* (Figure 7C, lower panel). The leading two Th2-specific enhancers activated under these conditions (number 4 and 8) were regulated positively by Th2 cytokines and negatively by T-bet. Similarly, of the leading Th1-specific enhancers that are inactivated, two were regulated by both T-bet and Th2 cytokines (numbers 5 and 7) while one was regulated primarily by T-bet (number 10). Hence, the forced downregulation of T-bet during reprogramming revealed strong quantitative T-bet dependence of the pivotal enhancers. Indeed, the transcriptome of *Tbx21^-/-^* Th1+2 cells was most similar to that of Th2 cells (Figure S7). The differential network of wildtype Th1 and Th2 cells revealed the same pivotal enhancer types (Figure 7C, upper panel) as between Th1 and *Tbx21^-/-^* Th1+2 cells, except for the mainly T-bet-dependent Th1-specific enhancer (number 10). Moreover, we found that in Th2 cells, an exclusively Th2-specific enhancer was active (number 9). This enhancer could not be switched on even during reprogramming of Th1 cells, suggesting that differentiated Th cell subsets are not perfectly interconvertible.

Analyzing multiple differential networks of different pairs of experimental conditions, we found enhancer types with strong T-bet dependence being key drivers of reprogramming upon loss of T-bet on one or both alleles (Figure 7D). However, under all reprogramming conditions enhancer types regulated by both Th2 cytokine conditions and T-bet dose were most important.

These finding support the notion that T-bet safeguards Th1 cell stability by modulating the enhancer landscape which is relevant upon subsequent activation of Th1 cells under adverse cytokine signals. Also the knockout of T-bet on both alleles under Th1→Th2 reprogramming reveals huge transcriptome similarity to Th2 cells though evading perfect interconvertibility.

## DISCUSSION

Understanding the topology and functional behavior of gene-regulatory networks is a key problem of genome biology, with applications to developmental biology (Davidson, 2010), immunology (Schulz et al., 2009; Spooner et al., 2009; Naldi et al., 2010), and many other fields. In the traditional view, the nodes of such networks are genes and the directed edges encode regulation of these genes by other gene products. The widespread use of logic functions (such as AND, OR etc.) to describe how two or more inputs to a gene are integrated (Davidson, 2010; Naldi et al., 2010) relies on the implicit assumption that the binding of multiple transcription factors realizes such logical functions. In higher organisms, and particularly in mammals, transcription factors regulate genes by binding to enhancers, and most genes are responsive to several enhancers at distinct genomic loci. Hence, the regulatory logic in mammalian gene-regulatory networks must build on the properties of individual enhancers and enhancer interactions with their target genes. Here, we introduce a new description of gene-regulatory networks that focuses on enhancers as elementary regulatory units. These enhancer-gene networks are described by bipartite graphs where enhancers and genes as nodes are connected by two types of edges: gene products (e.g., transcription factors) binding to enhancers and enhancers regulating their target genes. We introduce a computational method to infer such enhancer-gene networks from experimental data and apply it to understand the reprogramming of T helper cells.

A critical step of enhancer-gene network inference is the genome-wide mapping of enhancers to their target genes. Computationally, this has often been done in a much more simplified way by considering enhancers located in the upstream proximity of the gene of interest (Huang et al., 2016). More informed bioinformatic approaches integrate information on mapped enhancers and genomic contacts in three-dimensional space (Moore et al., 2020) or use chromatin features of enhancers in TADs (Kim et al., 2019). Here, we propose a different approach that utilizes multiple measurements of the same cell type under varying experimental conditions, causing activities of enhancers as well as expression of genes to change. We reasoned that co-variation of enhancer activity and gene expression indicates regulatory linkage (minimizing unwanted indirect effects by restricting enhancer-gene mappings to the same TAD and applying partial correlations to remove indirect effects within a TAD). A collection of well-characterized enhancers and their target genes in Th cells allowed us to learn a quantitative enhancer activity score from the experimental data that was based on graded values of three histone modifications: H3K4me1, H3K27ac, and H3K27me3. The first two of these are customarily used for identifying enhancers, yet H3K27me3 is not usually considered due to forming broad domains on chromatin. However, our data shows that the absence of H3K27me3 is critical for an active enhancer. Hence, we extend the previous classification of enhancers as either inactive (H3K4me1 without H3K27ac) or active (harboring H3K27ac) by a third, repressed, enhancer state if the enhancer is located in a H3K27me3 domain. It would be interesting to investigate whether the enhancer activity score could be refined by adding further molecular properties associated with enhancers, such as p300 binding.

A salient finding of enhancer analyses is that there are orders of magnitude more enhancers than genes in the genome, which appears to greatly complicate the inference of regulatory networks. For example, we found ~6000 enhancers for ~100 differentially expressed genes in Th1 and Th2 cells. The prediction of cytokine gene expression based on the activities of the respective enhancers indicates that multiple enhancers jointly control the transcript level. This could occur through simple addition (as suggested for *Ifng*) or cooperative action (possibly for *Il4*).

To dissect the complexity of the enhancer state space, we asked whether subsets of these enhancers share regulatory properties; we grouped enhancers with such shared properties in regulatory enhancer types. Encoding enhancer states in a ternary logic - active, inactive, and repressed - we found only approximately 10% of combinatorially possible types in our data (of which a subset of ~25% allowed reliable classification of Th1 and Th2 genes), suggesting that coherent, genome-wide modes of enhancer regulation govern the stability of Th1 cells and their reprogramming in the Th2 direction. Overall, these regulatory enhancer types fell into three categories: regulated by T-bet only, regulated by Th2 cytokines only, and regulated by both types of factors (mixed enhancer regulation). When analyzing genome-wide binding of key Th-cell transcription factors, we found that these three functionally identified categories are also characterized by distinct patterns of transcription factor binding. T-bet-regulated enhancers bound T-bet as well as STAT1 and STAT4. This is consistent with previous observations that these STAT factors open Th-cell-specific enhancers during Th1 cell differentiation (Vahedi et al., 2012). However, we also found strong T-bet binding, and the T-bet dependence was identified because enhancer activity in this regulatory type was gradually lost with declining *Tbx21* gene dose. Taken together, these findings suggest a scenario where enhancers are opened by STATs while enhancer activity is maintained by T-bet binding. This is akin to cell-type-specific enhancers in macrophages which are maintained by PU.1 binding (Natoli, 2010). Th2-cytokine-regulated enhancers predominantly bound STAT6 and GATA-3, while mixed enhancers bound all investigated factors.

It has been proposed that the cellular fate and identity of distinct cell types, as well as stability or plasticity in response to environmental challenges, rests on feedback architecture in gene-regulatory networks (Andrecut et al., 2011; Guantes and Poyatos, 2008). While previously such networks have been built by directly linking genes (e.g., a transcription factor to its target), our data allows the explicit consideration of enhancers. Given the coherent regulation of enhancer types by transcription factors, we chose enhancer types, rather than individual enhancers, as elementary building blocks (nodes), together with Th-cell-specific genes. A striking feature of the resulting enhancer-gene networks is their very high degree of connectivity, which sets them apart from conventional gene networks (Davidson, 2010). These connections form hundreds of positive feedback loops within the Th1- and Th2-specific gene clusters, which could self-reinforce the respective cell type identities. In previous work, negative regulatory interactions between Th1- and Th2-polarizing mechanisms have been prominently discussed (Löhning et al., 2002; Murphy and Reiner, 2002; Kanno et al., 2012). We also uncover such antagonistic interactions and make two key observations in this regard: First, negative interactions result when the presence of a transcription factor (e.g., T-bet) inactivates the respective enhancer (i.e., the transcription factor represses enhancer activity) whereas putative enhancer elements actively repressing gene expression did practically not occur. Second, such negative interactions were much rarer than positive feedback loops. The dominance of positive feedback compared with mutual repression suggests a mechanistic basis for the additive plasticity of Th-cell phenotypes in that attempted reprogramming does not abolish the starting phenotype (e.g., Th1) but rather adds new features, resulting, e.g., in Th1+2 cells.

The key feature of our analysis of enhancer-gene networks is the focus on regulatory enhancer types as nodes, rather than individual enhancers. If individual (distinct) enhancers were used, the network connectivity would be low (each enhancer has typically only one target gene) and regulatory similarity between genes could not be robustly inferred. By contrast, when using enhancer types, genes that are regulated similarly become strongly connected because they share enhancer types. Thus, Th1 and Th2 genes form clearly distinct clusters in force-directed network layouts (with only a small set of intermediate genes); we noticed that this key feature would be lost and Th1 and Th2 genes would become more intermingled in a classic gene network which does not consider enhancers. Enhancer types become crucial building blocks of the regulatory network. Importantly, our network constitutes a multi-digraph that retains basic mechanistic features in that gene interactions are always mediated by enhancers (leading, e.g., to gene (transcription factor)-enhancer-gene motifs); moreover, enhancer interactions are defined by shared transcription factors (constituting enhancer-gene-enhancer motifs).

Enhancer types provide a straightforward definition of condition-specific networks: If a given enhancer type is inactive in the considered condition then it is simply deleted from the network. Thus, formally the collection of all condition-specific enhancer-gene networks constitutes a multiplex network (Nicosia et al., 2013). In turn, the definition of condition-specific networks allowed us to identify key regulators that mediate the changes in cell phenotypes between conditions. To this end, we defined differential networks as the difference between two condition-specific networks, thus yielding the enhancer types that change activity. Moreover, we argue that enhancer types that are pivotal for a given transition, have high node importance (accounting for direct and indirect connectivity). This analysis revealed that enhancers regulated by both T-bet and Th2 cytokine input (mixed enhancer types) are most important for Th1 cell reprogramming, providing a general mechanism for plasticity in acquiring Th2 features combined with stability of Th1 features.

The real-time dynamics of the enhancer-gene networks introduced here pose entirely new, challenging questions. Dynamic models of conventional gene networks have been much studied and typically show two up to a handful of alternative steady states (bistability or multistability) that correspond to the specification of alternative cell fates. In stark contrast, the logic of epigenetic control implies that every individual enhancer could itself be a bistable switch (Dodd et al., 2007). The resulting extremely large state space of the network may allow for finely graded changes in enhancer activities and thus a quasi-continuum of stable cellular phenotypes. Such a mechanism could underlie the paradoxical observation that the molecular phenotypes of long-lived memory Th cells appear to lie on a continuum (Peine et al., 2013; Antebi et al., 2013; Huang, 2013).

## Author contributions

CK, QZ, ANH, ML, and TH designed the research. QZ and ANH performed the experiments. CK and QZ analyzed the data. CK, ANH, ML, and TH discussed the findings and their interpretation and visualization. CK and TH wrote the paper with input from ANH and ML.

## Acknowledgements

We thank Vivien Holecska and Isabel Panse for expert technical assistance and Roman M. Marek for the graphical design of the figures. This work was supported by the German Research Foundation (DFG-TRR241-A05 to ANH and grants LO 1542/4-1 and LO 1542/5-1 to ML), Volkswagen Foundation (Lichtenberg fellowships to ANH and ML), Willy Robert Pitzer Foundation (Pitzer Laboratory of Osteoarthritis Research to ML), Dr. Rolf M. Schwiete Foundation (Osteoarthritis Research Program to ML), and Berlin Institute of Health (Clinician Scientist grant to ANH).

## References

Afkarian, M., Sedy, J. R., Yang, J., Jacobson, N. G., Cereb, N., Yang, S. Y., Murphy, T. L. and Murphy, K. M. (2002). T-bet is a STAT1-induced regulator of IL-12R expression in naïve CD4+ T cells. Nat. Immunol. 3, 549–557.

Andersson, R. and Sandelin, A. (2019). Determinants of enhancer and promoter activities of regulatory elements. Nature Reviews Genetics 21, 71–87.

Andrecut, M., Halley, J. D., Winkler, D. A. and Huang, S. (2011). A general model for binary cell fate decision gene circuits with degeneracy: indeterminacy and switch behavior in the absence of cooperativity. PLoS ONE 6, e19358.

Ansel, K. M., Djuretic, I., Tanasa, B. and Rao, A. (2006). Regulation of Th2 differentiation and Il4 locus accessibility. Annu. Rev. Immunol. 24, 607–656.

Antebi, Y. E., Reich-Zeliger, S., Hart, Y., Mayo, A., Eizenberg, I., Rimer, J., Putheti, P., Pe’er, D. and Friedman, N. (2013). Mapping differentiation under mixed culture conditions reveals a tunable continuum of T cell fates. PLoS Biol. 11, e1001616.

Balasubramani, A., Mukasa, R., Hatton, R. D. and Weaver, C. T. (2010). Regulation of the Ifng locus in the context of T-lineage specification and plasticity. Immunol. Rev. 238, 216–232.

Barrington, C., Georgopoulou, D., Pezic, D., Varsally, W., Herrero, J. and Hadjur, S. (2019). Enhancer accessibility and CTCF occupancy underlie asymmetric TAD architecture and cell type specific genome topology. Nat Commun 10, 2908.

Battiston, F., Nicosia, V. and Latora, V. (2014). Structural measures for multiplex networks. Phys. Rev. E 89, 032804.

Brin, S. and Page, L. (1998). The anatomy of a large-scale hypertextual Web search engine. Computer Networks and ISDN Systems 30, 107–117.

Davidson, E. H. (2010). Emerging properties of animal gene regulatory networks. Nature 468, 911–920.

Dodd, I. B., Micheelsen, M. A., Sneppen, K. and Thon, G. (2007). Theoretical analysis of epigenetic cell memory by nucleosome modification. Cell 129, 813–822.

Ernst, J. and Kellis, M. (2010). Discovery and characterization of chromatin states for systematic annotation of the human genome. Nat. Biotechnol. 28, 817–825.

Geurts, P., Ernst, D. and Wehenkel, L. (2006). Extremely Randomized Trees. Machine Learning 36, 3–42.

Gong, Y., Lazaris, C., Sakellaropoulos, T., Lozano, A., Kambadur, P., Ntziachristos, P., Aifantis, I. and Tsirigos, A. (2018). Stratification of TAD boundaries reveals preferential insulation of super-enhancers by strong boundaries. Nat Commun 9, 542.

Gordon, S. and Martinez, F. O. (2010). Alternative Activation of Macrophages: Mechanism and Functions. Immunity 32, 593–604.

Guantes, R. and Poyatos, J. F. (2008). Multistable decision switches for flexible control of epigenetic differentiation. PLoS Comput. Biol. 4, e1000235.

Guo, W., Calixto, C. P. G., Tzioutziou, N., Lin, P., Waugh, R., Brown, J. W. S. and Zhang, R. (2017). Evaluation and improvement of the regulatory inference for large co-expression networks with limited sample size. BMC Syst Biol 11, 62.

Hegazy, A. N., Peine, C., Niesen, D., Panse, I., Vainshtein, Y., Kommer, C., Zhang, Q., Brunner, T. M., Peine, M., Fröhlich, A., Ishaque, N., Marek, R. M., Zhu, J., Höfer, T. and Löhning, M. (2022). Plasticity and lineage commitment of individual Th1 cells are determined by stable T-bet expression quantities. doi: 10.1101/2022.08.14.503916.

Hegazy, A. N., Peine, M., Helmstetter, C., Panse, I., Frhlich, A., Bergthaler, A., Flatz, L., Pinschewer, D. D., Radbruch, A. and Lhning, M. (2010). Interferons Direct Th2 Cell Reprogramming to Generate a Stable GATA-3+T-bet+ Cell Subset with Combined Th2 and Th1 Cell Functions. Immunity 32, 116–128.

Huang, J., Liu, X., Li, D., Shao, Z., Cao, H., Zhang, Y., Trompouki, E., Bowman, T. V., Zon, L. I., Yuan, G. C., Orkin, S. H. and Xu, J. (2016). Dynamic Control of Enhancer Repertoires Drives Lineage and Stage-Specific Transcription during Hematopoiesis. Dev. Cell 36, 9–23.

Huang, S. (2013). Hybrid T-helper cells: stabilizing the moderate center in a polarized system. PLoS Biol. 11, e1001632.

Kanno, Y., Vahedi, G., Hirahara, K., Singleton, K. and O’Shea, J. J. (2012). Transcriptional and epigenetic control of T helper cell specification: Molecular mechanisms underlying commitment and plasticity. Annu. Rev. Immunol. 30, 707–731.

Kim, D., An, H., Shearer, R. S., Sharif, M., Fan, C., Choi, J. O., Ryu, S. and Park, Y. (2019). A principled strategy for mapping enhancers to genes. Sci Rep 9, 11043.

Löhning, M., Richter, A. and Radbruch, A. (2002). Cytokine memory of T helper lymphocytes. Adv. Immunol. 80, 115–181.

Moore, J. E., Pratt, H. E., Purcaro, M. J. and Weng, Z. (2020). A curated benchmark of enhancer-gene interactions for evaluating enhancer-target gene prediction methods. Genome Biol. 21, 17.

Mullen, A. C., High, F. A., Hutchins, A. S., Lee, H. W., Villarino, A. V., Livingston, D. M., Kung, A. L., Cereb, N., Yao, T. P., Yang, S. Y. and Reiner, S. L. (2001). Role of T-bet in commitment of TH1 cells before IL-12-dependent selection. Science 292, 1907–1910.

Murphy, K. M. and Reiner, S. L. (2002). The lineage decisions of helper T cells. Nat. Rev. Immunol. 2, 933–944.

Nakayamada, S., Kanno, Y., Takahashi, H., Jankovic, D., Lu, K. T., Johnson, T. A., Sun, H. W., Vahedi, G., Hakim, O., Handon, R., Schwartzberg, P. L., Hager, G. L. and O’Shea, J. J. (2011). Early Th1 cell differentiation is marked by a Tfh cell-like transition. Immunity 35, 919–931.

Naldi, A., Carneiro, J., Chaouiya, C. and Thieffry, D. (2010). Diversity and plasticity of Th cell types predicted from regulatory network modelling. PLoS Comput. Biol. 6, e1000912.

Natoli, G. (2010). Maintaining cell identity through global control of genomic organization. Immunity 33, 12–24.

Nicosia, V., Bianconi, G., Latora, V. and Barthelemy, M. (2013). Growing Multiplex Networks. 111, 058701.

Ouyang, W., Lhning, M., Gao, Z., Assenmacher, M., Ranganath, S., Radbruch, A. and Murphy, K. M. (2000). Stat6-Independent GATA-3 Autoactivation Directs IL-4-Independent Th2 Development and Commitment. Immunity 12, 27–37.

Peine, M., Rausch, S., Helmstetter, C., Fröhlich, A., Hegazy, A. N., Kühl, A. A., Grevelding, C. G., Höfer, T., Hartmann, S. and Löhning, M. (2013). Stable T-bet(+)GATA-3(+) Th1/Th2 hybrid cells arise in vivo, can develop directly from naive precursors, and limit immunopathologic inflammation. PLoS Biol. 11, e1001633.

Reyes-Palomares, A., Gu, M., Grubert, F., Berest, I., Sa, S., Kasowski, M., Arnold, C., Shuai, M., Srivas, R. K., Miao, S., Li, D., Snyder, M. P., Rabinovitch, M. and Zaugg, J. B. (2020). Remodeling of active endothelial enhancers is associated with aberrant gene-regulatory networks in pulmonary arterial hypertension. Nature Communications 11.

Schulz, E. G., Mariani, L., Radbruch, A. and Höfer, T. (2009). Sequential polarization and imprinting of type 1 T helper lymphocytes by interferon-gamma and interleukin-12. Immunity 30, 673–683.

Spellberg, B. and Edwards, John E., J. (2001). Type 1/Type 2 Immunity in Infectious Diseases. Clinical Infectious Diseases 32, 76–102.

Spitz, F. and Furlong, E. E. M. (2012). Transcription factors: from enhancer binding to developmental control. Nature Reviews Genetics 13, 613–626.

Spooner, C. J., Cheng, J. X., Pujadas, E., Laslo, P. and Singh, H. (2009). A recurrent network involving the transcription factors PU.1 and Gfi1 orchestrates innate and adaptive immune cell fates. Immunity 31, 576–586.

Szabo, S. J., Kim, S. T., Costa, G. L., Zhang, X., Fathman, C. G. and Glimcher, L. H. (2000). A novel transcription factor, T-bet, directs Th1 lineage commitment. Cell 100, 655–669.

Vahedi, G., Takahashi, H., Nakayamada, S., Sun, H. W., Sartorelli, V., Kanno, Y. and O’Shea, J. J. (2012). STATs shape the active enhancer landscape of T cell populations. Cell 151, 981–993.

Wei, G., Abraham, B. J., Yagi, R., Jothi, R., Cui, K., Sharma, S., Narlikar, L., Northrup, D. L., Tang, Q., Paul, W. E., Zhu, J. and Zhao, K. (2011). Genome-wide analyses of transcription factor GATA3-mediated gene regulation in distinct T cell types. Immunity 35, 299–311.

Wei, L., Vahedi, G., Sun, H. W., Watford, W. T., Takatori, H., Ramos, H. L., Takahashi, H., Liang, J., Gutierrez-Cruz, G., Zang, C., Peng, W., O’Shea, J. J. and Kanno, Y. (2010). Discrete roles of STAT4 and STAT6 transcription factors in tuning epigenetic modifications and transcription during T helper cell differentiation. Immunity 32, 840–851.

Yamaguchi, T., Takizawa, F., Fischer, U. and Dijkstra, J. M. (2015). Along the Axis between Type 1 and Type 2 Immunity; Principles Conserved in Evolution from Fish to Mammals. Biology 4, 814–859.

Yue, F., Cheng, Y., Breschi, A., Vierstra, J., Wu, W., Ryba, T., Sandstrom, R., Ma, Z., Davis, C., Pope, B., Shen, Y., Pervouchine, D., Djebali, S., Thurman, R., Kaul, R., Rynes, E., Kirilusha, A., Marinov, G., Williams, B., Trout, D., Amrhein, H., Fisher-Aylor, K., Antoshechkin, I., DeSalvo, G., See, L., Fastuca, M., Drenkow, J., Zaleski, C., Dobin, A., Prieto, P., Lagarde, J., Bussotti, G., Tanzer, A., Denas, O., Li, K., Bender, M., Zhang, M., Byron, R., Groudine, M., McCleary, D., Pham, L., Ye, Z., Kuan, S., Edsall, L., Wu, Y., Rasmussen, M., Bansal, M., Kellis, M., Keller, C., Morrissey, C., Mishra, T., Jain, D., Dogan, N., Harris, R., Cayting, P., Kawli, T., Boyle, A., Euskirchen, G., Kundaje, A., Lin, S., Lin, Y., Jansen, C., Malladi, V., Cline, M., Erickson, D., Kirkup, V., Learned, K., Sloan, C., Rosenbloom, K., De Sousa, B., Beal, K., Pignatelli, M., Flicek, P., Lian, J., Kahveci, T., Lee, D., Kent, W., Santos, M., Herrero, J., Notredame, C., Johnson, A., Vong, S., Lee, K., Bates, D., Neri, F., Diegel, M., Canfield, T., Sabo, P., Wilken, M., Reh, T., Giste, E., Shafer, A., Kutyavin, T., Haugen, E., Dunn, D., Reynolds, A., Neph, S., Humbert, R., Hansen, R., De Bruijn, M., Selleri, L., Rudensky, A., Josefowicz, S., Samstein, R., Eichler, E., Orkin, S., Levasseur, D., Papayannopoulou, T., Chang, K., Skoultchi, A., Gosh, S., Disteche, C., Treuting, P., Wang, Y., Weiss, M., Blobel, G., Cao, X., Zhong, S., Wang, T., Good, P., Lowdon, R., Adams, L., Zhou, X., Pazin, M., Feingold, E., Wold, B., Taylor, J., Mortazavi, A., Weissman, S., Stamatoyannopoulos, J., Snyder, M., Guigo, R., Gingeras, T., Gilbert, D., Hardison, R., Beer, M. and Ren, B. (2014). A comparative encyclopedia of DNA elements in the mouse genome. Nature 515, 355–364.

