## Supplementary material for "Statistical Inference of Enhancer-Gene Networks Reveals Pivotal Role of T-bet Expression Intensity for T Helper Cell Fate"

### STAR METHODS

#### EXPERIMENTAL MODEL

##### Mice

C57BL/6 and C57BL/6 mice congenic for Thy1.1 (B6.PL-Thy1a/CyJ) mice were bred at the Institute of Experimental Medicine (FEM), Charité - University Medicine Berlin, and were used as recipients for adoptive cell transfers at the age of 8-12 wk. Tbx21$^{-/-}$ (Szabo et al., 2000) mice were purchased from Jackson laboratories (on a C57BL/6 background). SMARTA1 TCR-transgenic mice, which express a TCR specific for the LCMV epitope GP6180 were crossed to Thy1.1$^{+}$ B6.PL mice to generate TCRtg Thy1.1$^{+}$ mice.

##### Adoptive T cell transfer

Naïve LCMV-specific CD4$^{+}$ T cells ($1-3 \cdot 10^5$ cells) were given intravenously in 500 $\mu$l BSS to yield a seeding of approx. $1-3 \cdot 10^4$ cells per mouse (Hataye et al., 2006). In some experiments where the recipients were left uninfected, $0.5-1 \cdot 10^7$ cells were transferred to yield a seeding of approx. $0.5-1 \cdot 10^6$ cells per mouse.

##### Viruses

LCMV-ARM and LCMV-WE strains were propagated on BHK21 or L929 cells respectively. Mice were infected intravenously with 200 PFU.

##### T cell activation and differentiation

Naive MACS sorted LCMV-TCRtg CD4$^{+}$ T cells (CD4$^{+}$CD62L$^{+}$CD25$^{+}$) from spleens and lymph nodes isolated from the above mentioned mice were cultured in RPMI 1640 supplemented with 10% (vol/vol) FCS (Gibco), L-glutamine (2$\mu$M; Gibco), penicillin (100U/ml; Gibco), streptomycin (100$\mu$g/ml; Gibco) and b-mercaptoethanol (50nM; Sigma) in the presence of APCs and 1 $\mu$g/ml GP6480 peptide (Neosystem). For Th1 differentiation, 3 ng/ml IL-12 (R&D Systems) and 10 $\mu$g/ml anti-IL-4 (11B11) were added. For Th2 differentiation, 30 ng/ml IL-4 (Sigma-Aldrich) plus 10 $\mu$g/ml anti-IL-12 (C17.8) and 10 $\mu$g/ml anti-IFN-$\gamma$ (AN18.17.24) were used, as described previously (Hegazy et al., 2010). For in vitro conversion experiments, IL-25, IL-33, TSLP (Peprotech) were used at 10 ng/ml. Cell cultures were split on days 2 and 4. On day 6, cells were reactivated with fresh Thy1.2-depleted C57BL/6 splenocytes plus GP6480 peptide, 5 ng/ml IL-2 (R&D Systems), cytokines and anti-cytokine mAbs as above.

##### RNA Sequencing library preparation

RNA was extracted from $10^6$ cells using RNeasy mini kit (Qiagen) with a separate step for DNAse I digestion (Qiagen). ERCC Spike-in control mix-1 (Life technologies) was added to each sample prior to processing (1$\mu$L of a 1:10 dilution of mix-1). The rRNA depletion and library construction was processed with the ScriptSeq Complete Gold Kit (Epicentre) according to the manufacturers' instructions. The index for individual samples was added in order to multiplex 9 samples in each sequencing lane. The quality of libraries was assessed using a DNA1000 chip on the Bioanalyser (Agilent) and the concentration was measured using the Qubit DNA assay. Sequencing was performed on the Illumina platform with HiSeq 2000 paired-end 100 bp sequence type.

#### Chromatin Immunoprecipitation

ChIP experiments with Th cells were performed in biological duplicates using 106 cells. For fixation Th cells were incubated for 10 min at room temperature with 1% formaldehyde. Cross-linking was terminated by addition of glycine to a final concentration of 125 mM. Subsequently cells were incubated for 20 min at 4°C with a lysis buffer, nuclei were collected by centrifugation for 15 min at 20.000 g and 4°C and resuspended in a sonication buffer. The chromatin was fragmented by sonication (Covaris) to an average size of 150 bp. 1% of material was saved as "input". Antibodies were added to the sheared chromatin and DiaMag Protein A beads and incubated on a rotating wheel over night at 4°C. As negative controls rabbit and mouse normal IgG were used. The beads were washed four times with washing buffers. Immunocomplexes were disrupted by eluting 30 min at room temperature with an elution buffer. Eluates were reverse cross-linked by heating at 65°C for 4 h. Subsequently the DNA was purified with magnetic beads.

### METHOD DETAILS

#### Mapping and peak calling for ChIP-Seq data

Sequencing reads were aligned to the mouse genome (GRCm38.p4 build) using Bowtie (v1.2.1.1). Broad peak detection for ChIP-Seq data of histone modifications was performed by a local scanning window with an underlying Poissonian random read background model as implemented in SICER (v1.1). For fragment sizes of 150 bp we applied scanning window sizes of 200 bp and gap lengths of 600 bp for H3K4me1, H3K4me3, H3K27ac and H3K27me3 as well as gap lengths of 200 bp for H3K4me3. The background subtracted reads for each unit read bin of the length of the sliding window were then normalized w.r.t. library size according to

$$r_i' = \frac{r_i}{10^6 \cdot \sum_i r_i}$$

where $r_i$ denotes the unnormalized reads in some unit bin $i$. The result is given in reads per million (RPM).

#### Postprocessing of RNA-Seq data

Sequencing reads were aligned to the mouse genome (GRCm38.p4 build) using STAR (v2.4.0j) (Dobin et al., 2013). Read counts for each transcript were determined as the total number of reads mapped to unambiguous exons using htseq-count (v0.6.1) (Anders et al., 2015). For determining differentially upregulated genes between Th1 and Th2 conditions we used the function DESeq() from DESeq2 (v1.20) (Love et al., 2014). The genes were selected according to the multiple-testing-corrected p-value (according to Benjamini-Hochberg) and the largest logarithmic fold-change values (logFC). Additionally since we were interested in properly normalized transcript counts we performed a variance stabilizing transformation (VST) on the raw read counts using the function varianceStabilizingTransformation() from DESeq2.

#### Hidden Markov Model and model selection

Chromatin states were inferred genome-wide using ChromHMM (v1.10) (Ernst and Kellis, 2010). We trained a range from 5 to 25 states on the underlying 5 histone modifications from the ChIP-Seq data sets over all 8 experimental conditions. As noted in other publications (cite) we found that neither Bayesian Information Criterion (BIC) nor Akaike Information Criterion (AIC) scores were well discriminating approaches for model selection of Hidden Markov Models (HMM). Hence we facilitated the CompareModels routine of ChromHMM and determined a plateau in information gain between states of lower state models in comparison with the highest state model resulting in a 16 state model.

#### Parametrization of enhancer activity score

The learning sample of known enhancer elements was determined from various sources (Balasubramani et al., 2010; Jones and Flavell, 2005; Kanhere et al., 2012; Wilson et al., 2009; Yang et al., 2007) and is listed in Table 1.

| Gene | Enhancer | Element number |
|---|---|---|
| *Ifng* | CNS-54 | 8 |
| *Ifng* | CNS-6 | 11 |
| *Ifng* | Intron | 14 |
| *Ifng* | CNS+17-19 | 9 |
| *Ifng* | CNS+30 | 10 |
| *Ifng* | CNS+40 | 13 |
| *Ifng* | CNS+46 | 5 |
| *Ifng* | CNS+54 | 12 |
| *Tbx21* | -11.9 kb | 7 |
| *Tbx21* | -13.8 kb | 4 |
| *Il4* | HS1 | 2 |
| *Il4* | CNS2 | 3 |
| *Il10* | -9 kb | 1 |
| *Il10* | +6.45 kb | 6 |

Table 1: List of independently validated Th1 and Th2 enhancers used for parametrical learning of the enhancer activity score.

For optimization we employed the fminsearch() and fmincon() routines in MATLAB since we face a smooth non-linear optimization problem for random search initial conditions in the interval [-1, 1]. The optimization was then performed 1000 times for randomly sampled initial conditions. The obtained parameter values were stable with respect to initial conditions. We tested the validity of the parametrization with the MATLAB jackknife and bootstrap sampling functions jackknife() and bootstrp(). The latter was applied 1000 times. The corresponding $95\%$ confidence intervals were calculated via the quantile $z^*$ of a Gaussian distribution

$$z^* = 1 - \Phi^{-1}\left(1 - \frac{\alpha}{2}\right)$$

where $\Phi$ denotes the cumulative distribution function of a Gaussian and $\alpha$ as usual denotes the significance level. For the $95\%$ quantile, for a Gaussian being equivalent to a p-value of 0.05, we obtained the $95\%$ confidence interval by

$$\mathrm{CI}_{0.95} = \pm z^*_{0.975} \cdot \frac{\sigma}{\sqrt{n}}$$

with $n$ being the sample size. For a Gaussian distribution we have $z^*_{0.975} = 1.96$ leading to the resampling statistics

$$a_{\mathrm{CI}_{0.95}} = [1.2746, 1.3590] \qquad \sigma_{a_{\mathrm{CI}_{0.95}}} = [0.6316, 0.69128]$$

$$b_{\mathrm{CI}_{0.95}} = [-2.9634, -2.8197] \qquad \sigma_{b_{\mathrm{CI}_{0.95}}} = [1.0753, 1.1770]$$

for the bootstrap method.

#### Correlation Algorithm

The correlation method itself is implemented in a correlation algorithm specifically designed for this purpose. It builds upon the HMM segmentation as performed with ChromHMM (Ernst and Kellis, 2010). As the chromatin states are in some cases spatially very extended were segmented into smaller fragments down to a minimal resolution of 200 bp. Additionally an upper bound on the resolution was specified in order to circumvent extensive coarse-graining. The normalized histone modification reads $m_s$ in a resulting segment $s$ were obtained from its former bin values $m_{b_i}$ via

$$m_s = \sum_{i:|b_i \bigcap s| \neq 0} \frac{|b_i \bigcap s|}{b_i} \cdot m_{b_i} \,,$$

where the vertical bars denote the length of the segment. Additionally a statistical comparison between neighbouring segments was performed which led to a merging of similar segments. To this end the histone modification values $m_s$ within a segment of length $|s|$ were converted to densities

$$d_s = \frac{m_s \cdot 100}{|s|} \,.$$

For every segmented element with running index $i$ we now determined the tuple

$$\{\overline{\Delta d_s}, \sigma_{\Delta d_s}\}_{ij}$$

being the mean and the standard deviation of the difference of the histone modification densities for neighbouring segments $s_i$ and $s_j$ of all experimental input conditions $c$. The mean was obtained via

$$\overline{\Delta d_{s_{ij}}} = \frac{\sum_c (d_{s_{c,i}} - d_{s_{c,j}})}{c} \,.$$

In more detail segment $i$ is the left neighbour of segment $j$. We then defined boundaries within which neighbouring fragments have to lie in order to be merged. This could be specified as an additional input as well. To this end we chose a well documented gene locus to learn the mean and standard deviation distributions. We chose the *Ifng* locus to determine the learning sample distribution. From this we determined the inner 0.5-quantiles of the two distributions representing the median. Thus we assumed similarity of a total of half of the neighbouring segments within the training sample. From this we obtained the following boundaries on the similarity of neighbouring elements $ij$ for our correlation workflow based on the correlated histone modification segments:

$$\{\overline{\Delta d_s} \approx 0.025, \sigma_{\Delta d_s} \approx 0.05\}_{ij}\,.$$

Individual segments lying within these boundaries were iteratively merged. As soon as the merging condition was met the respective elements were being merged according to

$$d_s^* = \frac{d_{s_i} \cdot |s_i| + d_{s_{i-1}} \cdot |s_{i-1}|}{|s_i| + |s_{i-1}|}$$

Here again the vertical bars denote lengths of segments $s_i$. In effect we weighted each neighbour with its respective length and normalized the whole density in the end to its combined final length. This was repeated with the adjacent neighbours of the subsequently resulting element $d_s^*$ until the merging condition is not met anymore.

Eventually a lower cutoff for the obtained Pearson correlations of the merged segments was chosen to be 0.5 and a significance cutoff as $p < 0.1$. The correlation procedure was performed after specification of genes of interest on the respective topologically associating domains (TADs), which led to a limited amount of possible correlating segments in a sufficiently narrow region of interest. We furthermore only selected putative enhancer segments as determined by the HMM previously. For proper comparison we only considered experimental conditions with variations in Tbet dose for correlation.

#### Partial Correlations

Considering the pairwise correlation of a number of variables greater than two the mutual dependence of the variables on each other has to be disentangled first in order to remove spurious correlations of all other variables on any each pair. If one considers three variables $A$, $B$ and $C$ where a potential correlation between $A$ and $B$ is only mediated via the variable $C$ then the true partial (first-order) correlation coefficient between $A$ and $B$ given $C$ is given by

$$\rho_{AB|C} = \frac{\rho_{AB} - \rho_{AC}\rho_{BC}}{\sqrt{(1-\rho_{AC}^2)(1-\rho_{BC}^2)}}\,.$$

where $\rho_{ij}$ denote the ordinary zero-order correlation coefficients as for example in the case of Pearson or Spearman rank correlation. In the case of $N$ variables we have to determine the partial correlation of order $(N-2)$, which is defined recursively by lower order partial correlation coefficients:

$$\rho_{AB|C\cdots N} = \frac{\rho_{AB|D\cdots N} - \rho_{AC|D\cdots N}\rho_{BC|D\cdots N}}{\sqrt{(1-\rho^2_{AC|D\cdots N})(1-\rho^2_{BC|D\cdots N})}}\,.$$

In our case $A$ is the parametrized enhancer activity score, $B$ is the transcript to be correlated with and the vector $C \ldots N$ contains the other co-regulated transcripts for which we correct the correlation itself.

After having performed all partial correlations of a set of enhancers which was co-regulated according to the zero-order Pearson correlation with several transcripts we tested each partial correlation for statistical significance according to $p < 0.1$ and uniquely assigned the enhancer element to one transcript only if the difference between the highest and the second highest partial correlation of the element fulfilled $\Delta\rho > 0.2$.

#### Gene Expression Model

For a gene regulation model mediated by linear combinations of enhancer elements we assumed the following simple relation:

$$g_i = a \cdot \mathcal{H}_{ij} \cdot c_j + b\,,$$

with free parameters $a$ and $b$. Here $g_i$ denotes an element of the gene expression vector $\vec{g}$ with $i$ being the respective cell condition. Furthermore $c_j$ is an element of a correlation weight vector which assigns a weight to every enhancer segment around the gene under consideration depending on the resulting correlation value, which was inferred by the correlation algorithm. This is an assumption necessary to estimate the enhancer-specific deviations in the free parameter $a$, hence reducing the parameter space substantially. The length of index $j$ depends now on the number of independently called significantly correlating enhancer segments. The matrix with elements $\mathcal{H}_{ij}$ is a "conditions $\times$ enhancer segments" matrix containing the parametrized enhancer activity score values for each segment.

Furthermore we also investigated a *linear-exponential model* capturing potential cooperative behaviour of enhancer elements given by

$$g_i = e^{a\cdot\mathcal{H}_{ij}\cdot c_j + b}\,.$$

#### Extremely Randomized Trees and class-specific feature selection

##### Extremely Randomized Trees

We apply the Extremely Randomized Tree (ERT) method as introduced in (Geurts et al., 2006) and defined in the Python ExtraTreesClassifier() routine in the sklearn (v0.19.1) package on a weighted gene-enhancer-type matrix with a corresponding class vector. To this end we assigned each gene its specificity with respect to Th1 or Th2 differentiation decisions. As a learning sample we selected the set of 96 Th1- and Th2-specific gene transcripts and extracted enhancer types, denoted as $\mathscr{E}_j$ in the following, from the subset of significantly correlating segments. In addition to this we weighted every single enhancer instance $k$ with the width of the respective correlation segment denoted by $|s_k^*|$. Obviously the maximal number of instances $k$ could differ considerably from transcript to transcript leading to a matrix of segments $k_{ij}$ to be considered for each transcript $i$ and each enhancer type $j$. The resulting weighting coefficient is called $w_{jk}$. Every enhancer type is now treated as a feature or predictor variable for a certain set of gene transcripts. Hence we obtain the following sets of weighted class features

$$\left\{ \left( \sum_k w_{jk} \right)_{\mathscr{E}_j} \right\}_{\mathcal{G}}$$

around some gene transcript $\mathcal{G}$ belonging to gene class $\mathcal{C}$. Finally we obtained weighted transcript-feature matrix elements $\mathcal{M}_{ij}$, which read

$$\mathcal{M}_{ij} = \sum_{\mathbf{k}} w_{ij\mathbf{k}} = \begin{pmatrix} \sum_{\mathbf{k}}(w_{1,1})_{\mathbf{k}} & \sum_{\mathbf{k}}(w_{1,2})_{\mathbf{k}} & \cdots & \sum_{\mathbf{k}}(w_{1,j})_{\mathbf{k}} \\ \sum_{\mathbf{k}}(w_{2,1})_{\mathbf{k}} & \sum_{\mathbf{k}}(w_{2,2})_{\mathbf{k}} & \cdots & \sum_{\mathbf{k}}(w_{2,j})_{\mathbf{k}} \\ \vdots & \vdots & \ddots & \vdots \\ \sum_{\mathbf{k}}(w_{i,1})_{\mathbf{k}} & \sum_{\mathbf{k}}(w_{i,2})_{\mathbf{k}} & \cdots & \sum_{\mathbf{k}}(w_{i,j})_{\mathbf{k}} \end{pmatrix}$$

where $\mathbf{k} \equiv k_{ij}$ denotes the individual number of instances of each feature $j$ around transcript $i$. Furthermore the Gini impurity is used as a split estimator in the ERT method. Upon validation with a leave-one-out cross-validation method the obtained ranking stayed the same on average for the top-ranked features that exhibited the highest Gini impurity. In fact we recovered the top 10 feature ranks in all cases and the top 20 features with an accuracy of 93.44%.

##### Class-specific feature selection

The scoring algorithm we devised employed a novel intra-class specificity measure $\mathcal{I}_{\mathcal{C}}$ formally acting as a weight for the already determined Gini impurity $\mathcal{I}_{\text{Gini}}$. Hence we obtained a modified intra-class Gini impurity $\mathcal{I}^*_{\text{Gini}}$ which reads

$$\mathcal{I}^*_{\text{Gini}} = \mathcal{I}_{\mathcal{C}} \cdot \mathcal{I}_{\text{Gini}} \,.$$

The intra-class-specificity measure is now defined as

$$\mathcal{I}_{\mathcal{C},j} = \mathcal{N}_j \cdot \frac{\frac{\sum_i^{m \in \mathcal{C}} \mathcal{M}_{ij}(\mathcal{C})}{m_{\in \mathcal{C}}}}{\frac{\sum_{i,j}^{m,n \in \mathcal{C}} \mathcal{M}_{ij}(\mathcal{C})}{n_{\in \mathcal{C}}}} \cdot \frac{\frac{\sum_{i,j}^{m,n \notin \mathcal{C}} \mathcal{M}_{ij}(\neg \mathcal{C})}{n_{\notin \mathcal{C}}}}{\frac{\sum_i^{m \notin \mathcal{C}} \mathcal{M}_{ij}(\neg \mathcal{C})}{m_{\notin \mathcal{C}}}} \,.$$

Herein the index $j$ denotes the feature number with a maximum at $n$ for a certain class, $\mathcal{C}$ denotes the respective class (in this case either Th1 or Th2) and the maximal number of instances of a class is delimited by $m$. This normalization and averaging procedure takes all the weighted matrix entries from the feature matrix $\mathcal{M}_{ij}$ belonging to a certain class $\mathcal{C}$ and a certain feature – being a ternary enhancer type – and averages them for every feature separately. This is then normalized by the total weight of an average instance. The result additionally has to be normalized by the same measure for all other classes to make it independent of the total number of instances and classes. We also multiplied this by the number of instances where we found an entry larger than zero for each feature $j$ which we call $\mathcal{N}_j$. In the case of a binary class categorization this yielded two complimentary feature rankings for each class separately.

#### Network inference

##### Network architecture

We defined the set of network nodes $\mathscr{N}_i$ to consist of Th1- and Th2- specific genes as well as significantly correlating enhancer types being linked to the respective transcripts. The definition of the edges in the network is two-fold. First we defined directed edges originating from an enhancer type and ending at a gene. This means that direct edges between different enhancer types are forbidden leading to a bipartite network. The same also holds true for direct regulation between genes without enhancer mediation.

Also every transcript can be regulated by several correlated enhancer segments from the same enhancer type at the same time. We thus obtain a potential activation of a certain gene by multiple edges coming from one and the same enhancer type node. Such graphs are in general called *multigraphs*. In the particular case of directed edges such a graph is furthermore called *multidigraph* or *quiver*. Since we have to deal with a weighted multidigraph we treated the combination of edge weights between every two nodes by adding all available weights between a pairwise set of nodes $\{\mathscr{N}_i, \mathscr{N}_j\}$ for $i \neq j$ . As a consequence of this we defined the weighting $\mathcal{W}_{ij}$ of a directed *multi-edge* between a set of nodes $\{\mathscr{N}_i, \mathscr{N}_j\}$, where $i$ denotes an enhancer instance and $j$ denotes a gene transcript as

$$\mathcal{W}_{ij} = \sum_k w_{ijk} = \sum_k \rho(\mathcal{E}_{ik}, \mathcal{G}_j)$$

where $w_{ijk}$ denotes the $k$ individual edge weights going from $i \to j$ and $\rho(\mathcal{E}_{ik}, \mathcal{G}_j)$ is the Pearson correlation between an enhancer instance and the gene.

In order to obtain a bi-directional network we also define directed edges from transcription factors (TFs) to enhancer types. This is achieved through a weighting procedure of the ChIP-Seq binding data of TFs via the aforementioned ERT method, treating enhancer types as classes. The weighting in this case reads

$$\mathcal{W}_{ji} = \sum_k w_{jik} = \sum_k \mathcal{I}^*_{\text{Gini},jik} \,.$$

We note here that the individual modified Gini impurity $\mathcal{I}^*_{\text{Gini},ji}$ for a certain enhancer type with index $i$ was also normalized to one hence the individual edge weights $\mathcal{W}_{ij}$ and $\mathcal{W}_{ji}$ are comparable. We accounted for inhibiting edges with a negative edge weight if TFs acted as repressors. This was assessed by checking if the enhancer type exhibited a switch from an active to an inactive or repressed enhancer in a condition where the respective TF was binding.

The corresponding adjacency matrix for this weighted multidigraph with entries $\mathcal{A}_{ij}$ can be constructed in general from the edge weights $\mathcal{W}_{ij}$ and $\mathcal{W}_{ji}$ as follows:

$$\mathcal{A} = \begin{array}{l} \\ \text{TFs} \\ \text{Enhancer types} \end{array} \begin{array}{c} \begin{array}{cc} \text{Genes} & \text{Enhancer types} \end{array} \\ \begin{pmatrix} 0 & \mathcal{W}_{ji} \\ \mathcal{W}_{ij} & 0 \end{pmatrix} \end{array}$$

Networks are visualized via a force-directed graph drawing using Cytoscape (v3.4.0). The nodes of the conditions-specific networks were kept fixed with respect to their force-directed positions of the full network.

#### Multiplex and differential networks

Multiplex networks are condition-specific networks that may have common overlaps between certain nodes and edges. We assume that as soon as a certain enhancer-type loses its enhancer activity in a certain condition, then the node is removed from this particular multiplex network. This also leads to the loss of the associated edges and changes the connectivity as well as the topology of the network. Furthermore genes are depicted as switched off (white) in a certain condition as soon as their VST-normalized expression value is $g < 0,5 \cdot g_{\text{max}}$ in that condition.

Differential networks constitute the pairwise difference of the respective multiplex networks leading to up- and downregulated edges and including nodes that are added as well as removed between the respective conditions.

#### Node importance ranking

The importance ranking of nodes as facilitated in the differential networks uses the same method as applied in the PageRank algorithm (Brin and Page, 1998). To this end we make use of the function pagerank() from the Python networkx-package (v1.11). This means creating a one-step raw transition-matrix

$$\mathcal{P}_{ij} = k_{\text{out},i}^{-1} \cdot |\mathcal{W}_{ij}| \tag{1}$$

transitioning from node $i$ to node $j$ via a probability being determined by the absolute weight between those nodes and being normalized by the weighted out-degree of node $i$. A random walk of $N$ steps starting at $i$ and ending at $j$ is now obtained by $N$ multiplications of the transition matrix with itself, hence computing $\mathcal{P}_{ij}^N$. More precisely this defines a time-discrete time-forward random walk constituting a Markov chain over $N$ steps. As stated in the *Perron-Frobenius theorem for ergodic Markov chains* one can after appropriate transformations of the one-step raw transition-matrix to a stochastic irreducible matrix obtain a largest eigenvalue of the matrix with $\lambda_1 = 1$ . Accordingly the characteristic equation yields

$$\pi_j^T \mathcal{P}_{ij} = \pi_j^T \ . \tag{2}$$

with left eigenvector $\pi^T$. This eigenvector constitutes a stationary distribution ranking all network nodes time-independently w.r.t. to their respective stochastic importance within the network .

### References

Anders, S., Pyl, P. T. and Huber, W. (2015). HTSeq–a Python framework to work with high-throughput sequencing data. Bioinformatics *31*, 166–169.

Balasubramani, A., Mukasa, R., Hatton, R. D. and Weaver, C. T. (2010). Regulation of the Ifng locus in the context of T-lineage specification and plasticity. Immunol. Rev. *238*, 216–232.

Brin, S. and Page, L. (1998). The anatomy of a large-scale hypertextual Web search engine. Computer Networks and ISDN Systems *30*, 107–117.

Dobin, A., Davis, C. A., Schlesinger, F., Drenkow, J., Zaleski, C., Jha, S., Batut, P., Chaisson, M. and Gingeras, T. R. (2013). STAR: ultrafast universal RNA-seq aligner. Bioinformatics *29*, 15–21.

Ernst, J. and Kellis, M. (2010). Discovery and characterization of chromatin states for systematic annotation of the human genome. Nat. Biotechnol. *28*, 817–825.

Geurts, P., Ernst, D. and Wehenkel, L. (2006). Extremely Randomized Trees. Machine Learning *36*, 3–42.

Hataye, J., Moon, J. J., Khoruts, A., Reilly, C. and Jenkins, M. K. (2006). Naïve and Memory $CD4^+$ T Cell Survival Controlled by Clonal Abundance. Science *312*, 114–116.

Hegazy, A. N., Peine, M., Helmstetter, C., Panse, I., Frhlich, A., Bergthaler, A., Flatz, L., Pinschewer, D. D., Radbruch, A. and Lhning, M. (2010). Interferons Direct Th2 Cell Reprogramming to Generate a Stable GATA-3+T-bet+ Cell Subset with Combined Th2 and Th1 Cell Functions. Immunity *32*, 116–128.

Jones, E. A. and Flavell, R. A. (2005). Distal enhancer elements transcribe intergenic RNA in the IL-10 family gene cluster. J. Immunol. *175*, 7437–7446.

Kanhere, A., Hertweck, A., Bhatia, U., Gökmen, M. R., Perucha, E., Jackson, I., Lord, G. M. and Jenner, R. G. (2012). T-bet and GATA3 orchestrate Th1 and Th2 differentiation through lineage-specific targeting of distal regulatory elements. Nat Commun *3*, 1268.

Love, M. I., Huber, W. and Anders, S. (2014). Moderated estimation of fold change and dispersion for RNA-seq data with DESeq2. Genome Biol. *15*, 550.

Nakayamada, S., Kanno, Y., Takahashi, H., Jankovic, D., Lu, K. T., Johnson, T. A., Sun, H. W., Vahedi, G., Hakim, O., Handon, R., Schwartzberg, P. L., Hager, G. L. and O'Shea, J. J. (2011). Early Th1 cell differentiation is marked by a Tfh cell-like transition. Immunity *35*, 919–931.

Szabo, S. J., Kim, S. T., Costa, G. L., Zhang, X., Fathman, C. G. and Glimcher, L. H. (2000). A novel transcription factor, T-bet, directs Th1 lineage commitment. Cell *100*, 655–669.

Vahedi, G., Takahashi, H., Nakayamada, S., Sun, H. W., Sartorelli, V., Kanno, Y. and O'Shea, J. J. (2012). STATs shape the active enhancer landscape of T cell populations. Cell *151*, 981–993.

Wei, G., Abraham, B. J., Yagi, R., Jothi, R., Cui, K., Sharma, S., Narlikar, L., Northrup, D. L., Tang, Q., Paul, W. E., Zhu, J. and Zhao, K. (2011). Genome-wide analyses of transcription factor GATA3-mediated gene regulation in distinct T cell types. Immunity *35*, 299–311.

Wilson, C. B., Rowell, E. and Sekimata, M. (2009). Epigenetic control of T-helper-cell differentiation. Nat. Rev. Immunol. *9*, 91–105.

Yang, Y., Ochando, J. C., Bromberg, J. S. and Ding, Y. (2007). Identification of a distant T-bet enhancer responsive to IL-12/Stat4 and IFNgamma/Stat1 signals. Blood *110*, 2494–2500.

### SUPPLEMENTAL INFORMATION

| Th1 transcript | Th2 transcript |
|---|---|
| Ifng-201 | A430108G06Rik-002 |
| Tbx21-001 | Gm12214-001 |
| Runx3-001 | Gm17334-201 |
| Runx3-002 | Gm22275-201 |
| Cxcr3-001 | Il4-001 |
| Il2-001 | Il4-003 |
| Il12rb2-001 | Il5-001 |
| Il12rb2-002 | Il13-001 |
| Eomes-001 | Irf1-002 |
| Eomes-003 | Irf1-008 |
| Ccl5-001 | Kif3a-001 |
| Klri2-001 | Kif3a-005 |
| Itga1-201 | Rad50-001 |
| Klrc1-001 | Sept8-002 |
| Klrc1-002 | Sept8-004 |
| Klrc1-201 | Gata3-001 |
| Klrc1-202 | Il10-001 |
| Klrc2-201 | Ccr4-201 |
| Klrc2-001 | Ccr1-201 |
| Klrc2-002 | Areg-201 |
| Klrc2-004 | Pparg-202 |
| Stat1-001 | Pparg-201 |
| Stat1-004 | Il9r-004 |
| Stat1-006 | Il9r-003 |
| Stat1-007 | Il9r-001 |
| Stat1-008 | Asb2-002 |
| Stat1-009 | Asb2-001 |
| Stat4-001 | Lrrc32-201 |
| Stat4-002 | Adamtsl3-001 |
| Il18r1-202 | Adamtsl3-005 |
| Il18rap-001 | Gja1-201 |
| Ccr2-001 | Treml2-201 |
| Ccr5-001 | Sell-001 |
| Fasl-001 | Sell-002 |
| Fasl-002 | Sell-003 |
| Smpdl3b-001 | Stat6-001 |
| Klrg1-001 | Il1rl1-001 |
| Kcnj8-201 | Il1rl1-002 |
| Gldc-201 | Il1rl1-003 |
| Ly6c2-001 | Ifngr2-001 |
| Clec12a-001 | Chil3-001 |
| Dpysl3-001 | Inpp4b-008 |
| Galnt3-001 | Mctp1-001 |
| Klre1-201 | Efna5-001 |
| Exph5-001 | Igfbp4-001 |
| Klrb1c-003 | Slc4a4-001 |
| | Bhlhe41-001 |
| | St8sia6-001 |
| | Cd83-201 |
| | Cyp11a1-001 |

**Table S1**: List of 46 Th1 and 50 Th2 transcripts as determined in (Wei et al., 2011).

**S1: Experimental plan demonstrating the adoptive transfer procedure and culture conditions**

Naïve WT, Tbx21$^{+/-}$ and Tbx21$^{-/-}$ LCMV specific CD4$^{+}$ Thy1.1$^{+}$ cells were adoptively transferred into C57BL/6 mice. Recipient mice were infected with LCMV. 10 days after infection, CD4$^{+}$ Thy1.1$^{+}$ donor T cells were isolated and re-activated for 2 rounds with weekly re-activation under neutral or Th2 conditions. Subsequently, *in vitro*-re-activated CD4$^{+}$ Thy1.1$^{+}$ cells were re-transferred into naïve C57BL/6 mice. For all of the cells RNA-Seq was performed as well as ChIP-Seq for H3K27ac, H3K4me3, H3K4me1, H3K9me3, and H3K27me3.

**S2: Histone modification peak data for both experimental replicates and all experimental conditions**

Significant histone modification peaks for 3K27ac, H3K4me3, H3K4me1, H3K9me3, and H3K27me3 at the signature Th1 cytokine locus of Ifng for all experimental cell conditions. On the left side we listed all permissive marks for both replicates of all experimental conditions; on the right hand side we listed the same for the repressive marks.

**S3: Emission parameter correlation for the Hidden Markov model**

Heatmap of correlation values (shades of grey) between emission parameters between Hidden Markov Models with varying state numbers. The y-axis depicts all hidden states of the 25-state model which was considered to be the largest model for the observed five histone modifications. The x-axis shows all models that were trained from 5 up to 25 states. The heatmap analyses the overall correlation between each state from the 25-state model with each trained model. A drop in correlation when going from higher to lower state models on the x-axis for one state uncovers an information loss with the removal of that particular state.

**S4: Detailed enhancer-gene correlation analysis at Ifng**

(A) Individual Pearson correlations for the enhancer learning sample (numbering see Table S1) of H3K4me1, H3K27ac, H3K27me3 with the respective gene transcript abundances. Additionally we show the chosen Pearson correlation of the enhancer activity score as well as its respective Spearman rank correlation. The distributions of the respective correlations are shown on the right.

(B) Bootstrapping distributions of the parameters $a$ and $b$ from the enhancer activity score for H3K27ac and H3K27me3 respectively.

**S5: Chromatin states and significant correlations at the Tbx21 superenhancer locus**

Tbx21 locus with the according HMM state classification for different experimental conditions. The location of the superenhancer is shown as well as the predicted significant correlations from the enhancer activity score in addition to binding sites of Tbet and p300 from experimental ChIP-Seq data sets (Nakayamada et al., 2011; Vahedi et al., 2012).

**S6: Full inferred enhancer-gene network**

Full enhancer-gene network according to a force-directed depiction. Red nodes indicate Th2 specificity, blue nodes indicate Th1 specificity, whereas black enhancer nodes (circles) are statistically cell-type-unspecific. Black network edges indicate activation, (sparse) red network edges indicate inactivation.

**S7: Transcriptome distinction of experimental conditions**

Visualization of the similarities within the full transcriptome information of the considered experimental conditions and their replicates. The analysis is provided by a principal component analysis (PCA) showing the leading principle components (PC1 and PC2) accounting for 48% and 25% of the variance respectively. The PC1 is able to distinguish nicely between naïve, Th1 and Th1→Th2 reprogramming conditions. The PC2 on the other hand focusses on the role of T-bet by exhibiting equal displacements from the neutral to the heterozygous and the knock-out conditions for both Th1 and Th1→Th2 cells. Also the transcriptome similarity between KO/2 and Th2 becomes quite evident as also mirrored by the differential network analyses visualized in Figure 7.

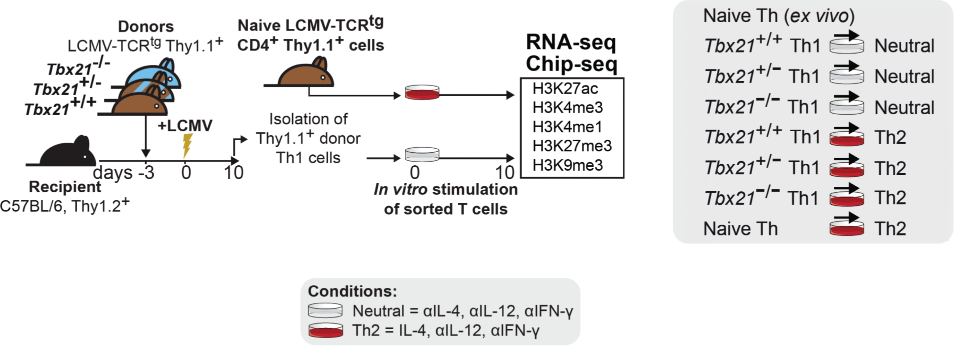
Donors
LCMV-TCRtg Thy1.1+
Tbx21-/-
Tbx21+/-
Tbx21+/+
+LCMV
Recipient
C57BL/6, Thy1.2+
days -3 0 10
Naive LCMV-TCRtg CD4+ Thy1.1+ cells
Isolation of Thy1.1+ donor Th1 cells
0 10
In vitro stimulation of sorted T cells
RNA-seq
Chip-seq
H3K27ac
H3K4me3
H3K4me1
H3K27me3
H3K9me3
Naive Th (ex vivo)
Tbx21+/+ Th1 Neutral
Tbx21+/− Th1 Neutral
Tbx21−/− Th1 Neutral
Tbx21+/+ Th1 Th2
Tbx21+/− Th1 Th2
Tbx21−/− Th1 Th2
Naive Th Th2
Conditions:
Neutral = αIL-4, αIL-12, αIFN-γ
Th2 = IL-4, αIL-12, αIFN-γ

Fig. S1

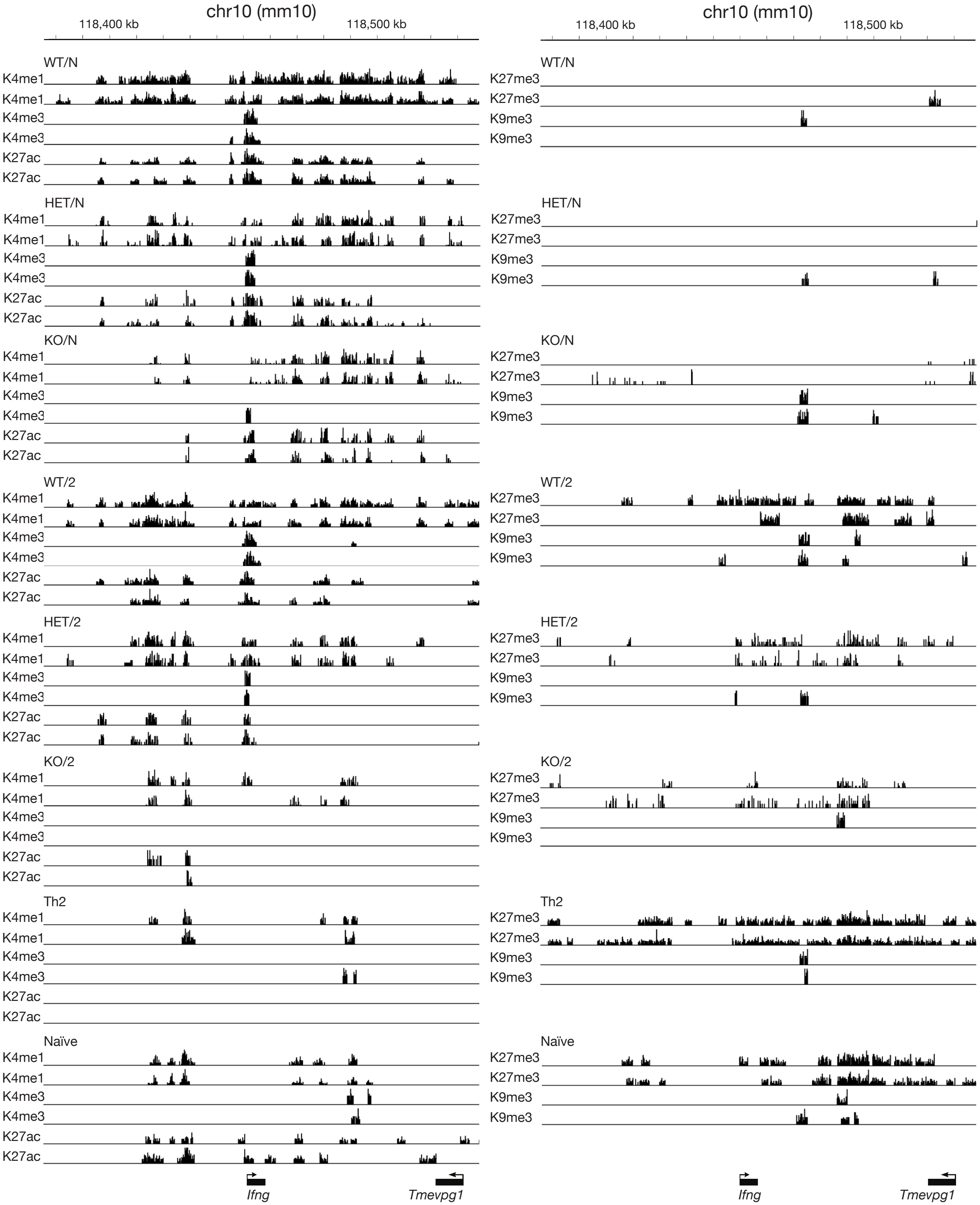

chr10 (mm10)
118,400 kb
118,500 kb
WT/N
K4me1
K4me3
K27ac
K27me3
K9me3
HET/N
KO/N
WT/2
HET/2
KO/2
Th2
Naïve
Ifng
Tmevpg1

Fig. S2

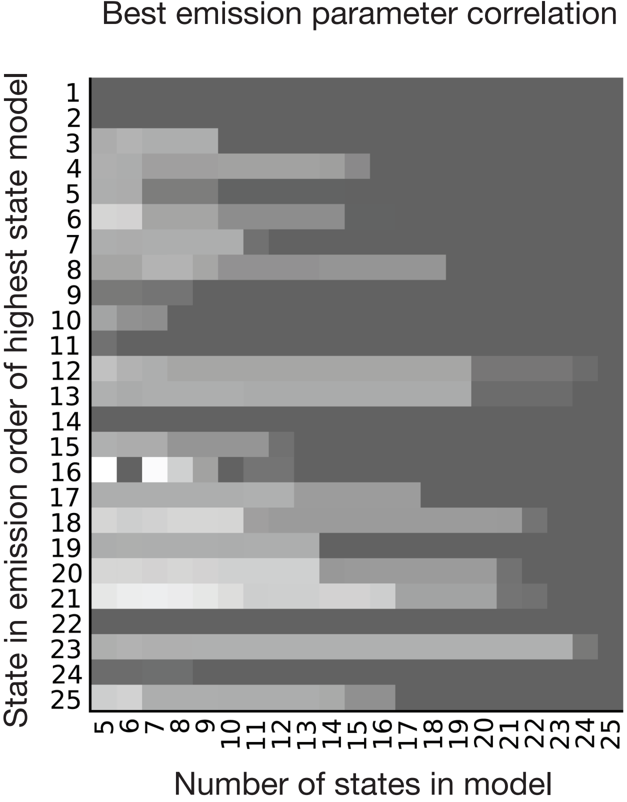
Best emission parameter correlation
State in emission order of highest state model
Number of states in model

Fig. S3

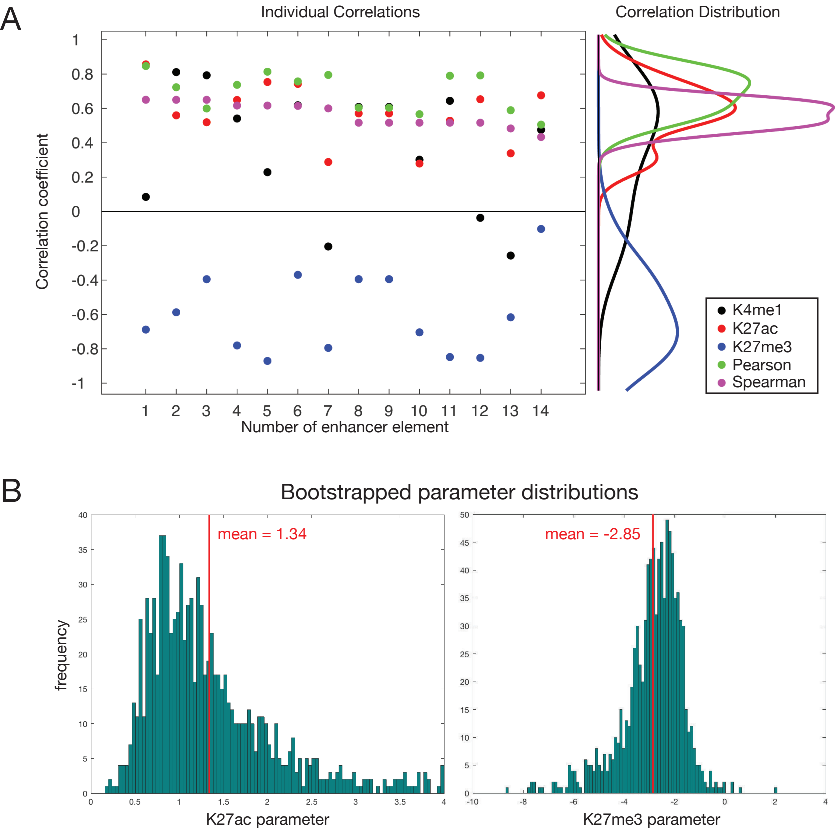
A
Individual Correlations
Correlation Distribution
Correlation coefficient
Number of enhancer element
K4me1
K27ac
K27me3
Pearson
Spearman
B
Bootstrapped parameter distributions
frequency
mean = 1.34
mean = -2.85
K27ac parameter
K27me3 parameter

Fig. S4

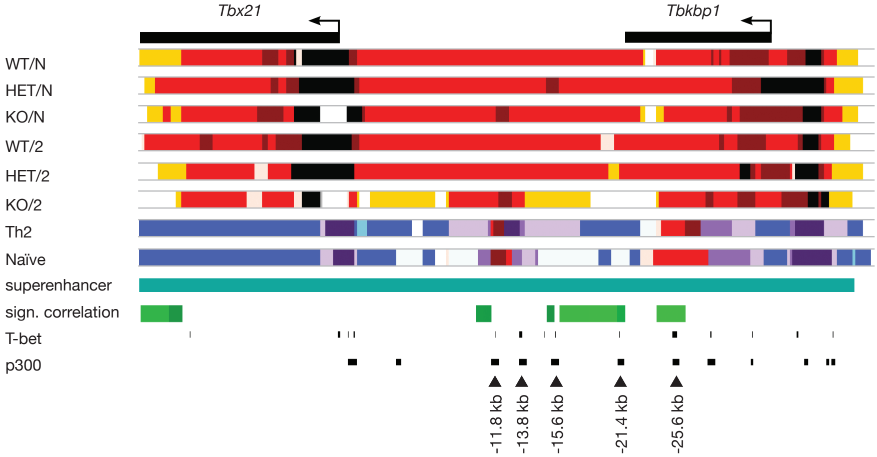
Tbx21
Tbkbp1
WT/N
HET/N
KO/N
WT/2
HET/2
KO/2
Th2
Naïve
superenhancer
sign. correlation
T-bet
p300
-11.8 kb
-13.8 kb
-15.6 kb
-21.4 kb
-25.6 kb

Fig. S5

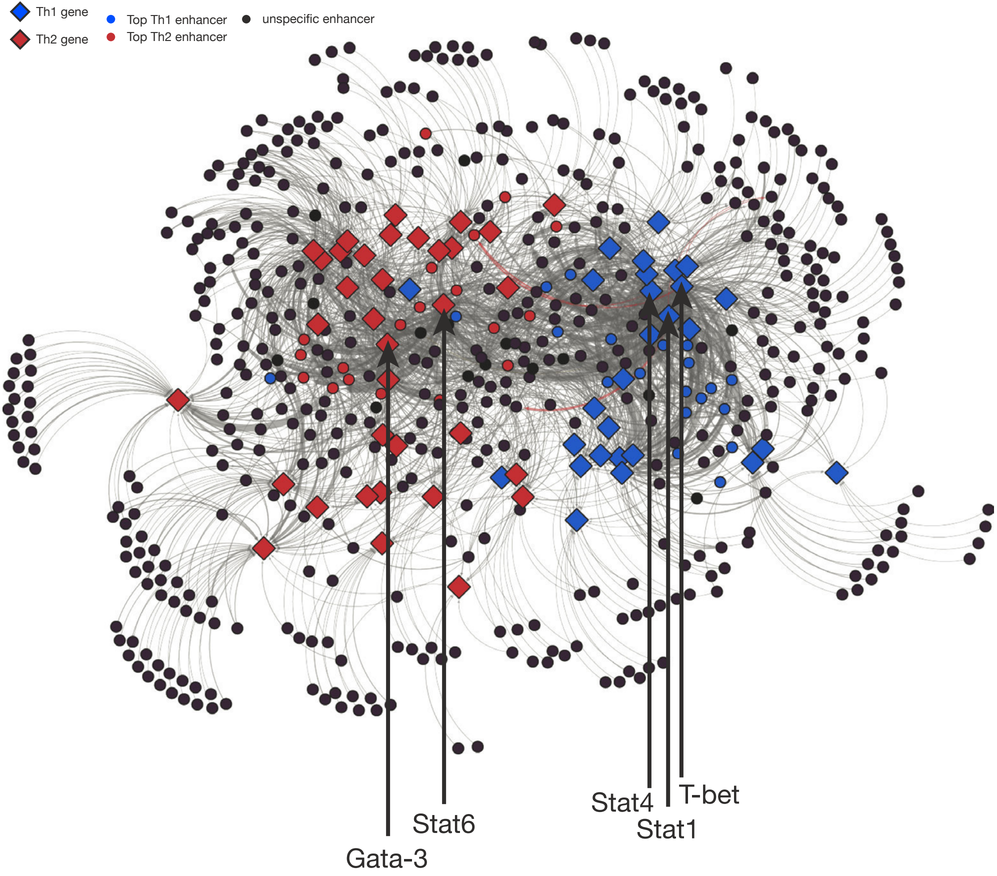
Th1 gene
Th2 gene
Top Th1 enhancer
Top Th2 enhancer
unspecific enhancer
Gata-3
Stat6
Stat4
Stat1
T-bet

Fig. S6

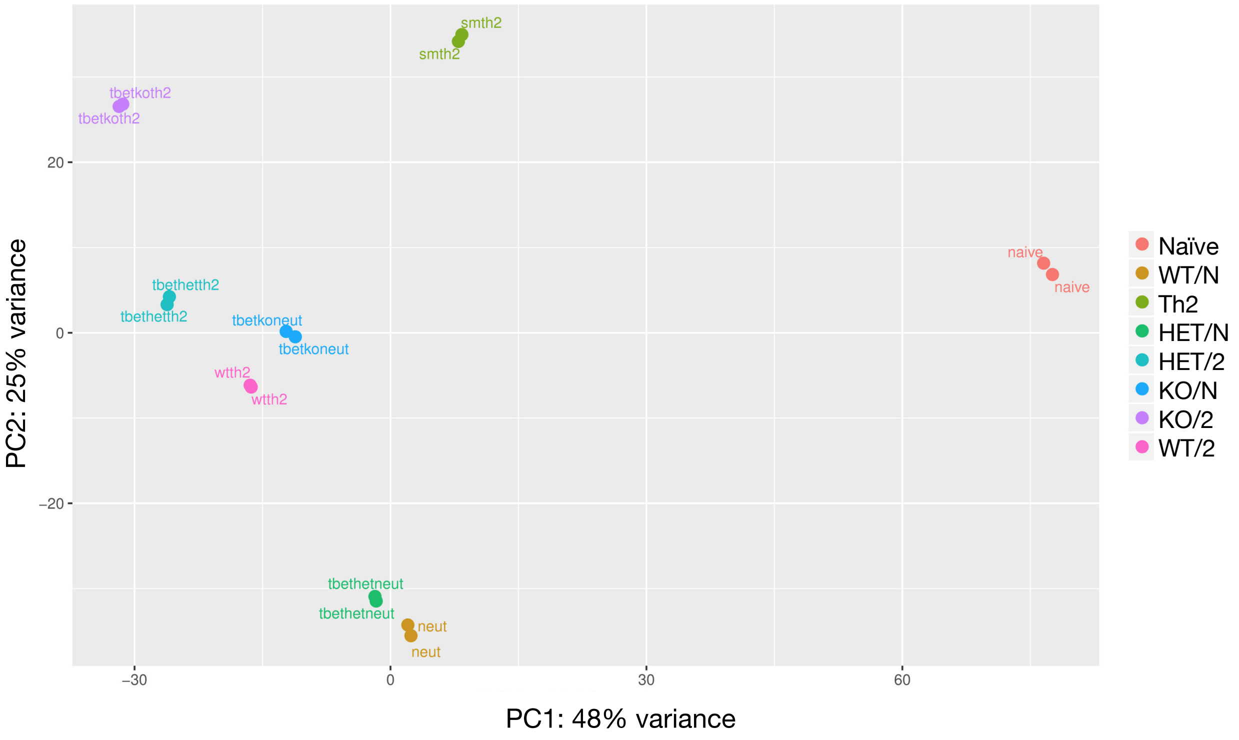

smth2
smth2
tbetkoth2
tbetkoth2
naive
naive
tbethetth2
tbethetth2
tbetkoneut
tbetkoneut
wtth2
wtth2
tbethetneut
tbethetneut
neut
neut
20
0
−20
−30
0
30
60
PC2: 25% variance
PC1: 48% variance
Naïve
WT/N
Th2
HET/N
HET/2
KO/N
KO/2
WT/2

Fig. S7
